# Cahn-Hilliard dynamical models for condensed biomolecular systems

**DOI:** 10.1101/2025.07.13.664571

**Authors:** Sarah M. Groves, Min-Jhe Lu, Astrid Catalina Alvarez-Yela, Monserrat Gerardo-Ramírez, P. Todd Stukenberg, John S. Lowengrub, Kevin A. Janes

## Abstract

Biomolecular condensates create dynamic subcellular compartments that alter systems-level properties of the networks surrounding them. One model combining soluble and condensed states is the Cahn-Hilliard equation, which specifies a diffuse interface between the two phases. Customized approaches required to solve this equation are largely inaccessible. Using two complementary numerical strategies, we built stable, self-consistent Cahn-Hilliard solvers in Python, MATLAB, and Julia. The algorithms simulated the complete time evolution of condensed droplets as they dissolved or persisted, relating critical droplet size to a coefficient for the diffuse interface in the Cahn-Hilliard equation. We applied this universal relationship to the chromosomal passenger complex, a multi-protein assembly that reportedly condenses on mitotic chromosomes. The fully constrained Cahn-Hilliard simulations yielded dewetting and coarsening behaviors that closely mirrored experiments in different cell types. The Cahn-Hilliard equation tests whether condensate dynamics behave as a phase-separated liquid, and its numerical solutions advance generalized modeling of biomolecular systems.

## INTRODUCTION

Phase separation arises whenever random-walk diffusion is countered by attractive forces that are strong enough to offset the energetic penalty of forming an interface between phases. In biology, such attractive forces may be biomolecular,^1,2^ adhesive,^3,4^ mechanical,^4–6^ or unknown.^7^ Phase-separating behavior has been documented at biological length scales ranging from subdomains of organellar membranes^8^ to entire ecological communities.^9^ One area of interest for systems biology is the condensation of biomolecules as 0.1–1 µm droplets within cells, which are thought to change the emergent properties of networks by insulation,^10^ kinetic modulation,^11^ and buffering.^12^

Condensed two-phase systems can be modeled as a free-boundary problem that posits a zero-thickness interface with a surface tension and accompanying force balance. However, this approach is problematic at small length scales, which must be considered when very small droplets form, as in the Rayleigh instability,^13,14^ or dissolve, as in Ostwald ripening.^15^ An alternative that dates back to Poisson is to consider a diffuse boundary, which changes between phases over a length scale that emerges from an imbalance in intermolecular forces.^16^ An order parameter identifies one of the chemical states (Box 1), and governing equations follow an energy variational approach. One classic example is the Cahn-Hilliard equation,^17^ which evolves by diffusion arising from the variational derivative of a free energy. The Cahn-Hilliard free energy contains a local function with two minima (one for each chemical state; Box 1) and a gradient term that penalizes the formation of interfacial regions (Box 2). The Cahn-Hilliard model is appealing for two-phase chemical transport because it conserves mass and dissipates energy toward a thermodynamic minimum. Indeed, many of the biological examples cited above have been modeled as Cahn-Hilliard systems.

For most two-phase phenomena, there is not an exact solution for the Cahn-Hilliard equation.

Numerical approximations are possible, but they require specialized approaches to address space/time-step restrictions, local nonlinearities, and nonconvexity. Published algorithms demonstrate proof of concept but historically do not include freely available code for redeployment. Rare exceptions^18–20^ are limited to specific boundary conditions that may not always be biologically appropriate. A general toolkit for Cahn-Hilliard models may facilitate its application to other examples of biological condensation.^7^

Toward modeling the dynamics of biomolecular condensates within cells,^21,22^ here we assemble and validate a multi-language suite^23^ of numerical methods for solving the Cahn-Hilliard equation. The algorithms are stable, self-consistent, and computationally suitable for spatial meshes of up to 2^9^ × 2^9^= 262,144 elements. Applying the methods to hundreds of long-term, high-resolution Cahn-Hilliard simulations, we define the critical equilibrium radius (*R*_*eq*_) for a stable, condensed droplet across a tenfold range of plausible diffuse interfaces. Focusing on a protein complex that reportedly forms biomolecular condensates when bound to mitotic chromosomes,^24^ we estimate *R*_*eq*_ and with the corresponding diffuse interface simulate different recruitment patterns as a Cahn-Hilliard process. Although the phase-separating behavior of this complex is debated,^25^ we find that its dewetting and ripening characteristics are remarkably consistent with theory. As a Method, this work provides access to reliable tools for abstracting dynamic interfaces in biological systems.

### Box 1. Chemical states and energy functions in a Cahn-Hilliard system

The Cahn-Hilliard equation models the dynamics of dimensionless chemical states. It is straightforward to convert a real chemical concentration (*rcc* [=] mass/length^3^) to a dimensionless relative chemical state (*c* [=] dimensionless) by normalizing to the concentration of the species at saturation (*rcc*_*max*_ [=] mass/length^3^): *c* = *rcc* / *rcc*_*max*_. In a purely diffusive system, the free energy (*F*) of *c* from Flory–Huggins solution theory is approximated by a parabola^26^ with a minimum at the well-mixed average (*c*_*avg*_): *F*(*c*) = *α*(*c* − *c*_*avg*_)^2^ + β. *α* and *β* specify the units and reference state for *F*, but they do not impact the equilibrium solution of minimum free energy (and thus zero chemical potential) where *dF*/*dc* = 0 (Figure B1A, yellow). To simplify the presentation and help with later scaling arguments, we keep *F* dimensionless by normalizing *F* by *k*_*B*_*T* (where *k*_*B*_ is the Boltzmann constant and *T* is absolute temperature) and take *α* = 1, *β* = 0.

For a Cahn-Hilliard two-phase system,^17^ an alternative double-well free energy function is used to approximate a Flory–Huggins solution with interacting phases (also known as the regular solution theory; Figure B1A, black), which we begin to introduce here in terms of stable minima at *c* = 0, 1 and *α* = 1, *β* = 0 (Figure B1A, green):

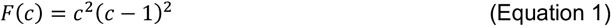

Note that any pair of stable minima can be rescaled to fall on a [0,1] interval by linearly shifting the minimum and dividing by the range. For stable minima *γ* = *γ*_*–*_, *γ*_*+*_:

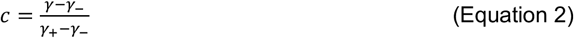

For mathematical convenience, it is advantageous to convert the double-well function of Equation 1 to a pair of qualitative chemical states at *ϕ* = –1, 1 by linearly translating and scaling (Figure B1B, purple):

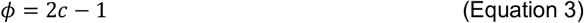

to yield:

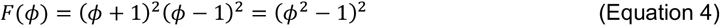

This symmetric double-well function of Figure B1B is found most often in the Cahn-Hilliard literature, but system behavior is alternatively described according to its spinodal point—the minimum concentration at which two phases stably coexist. The spinodal point is equivalent to the leftmost inflection point of the free energy. Taking Equation 4 as an equality, differentiating twice, and solving for the negative root yields:

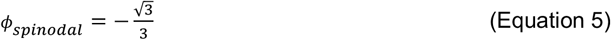

The spinodal point generalizes to any concentration by taking Equation 5, converting *ϕ* to *c* and then *c* to *γ* (Equations 2 and 3; Figure B1C):

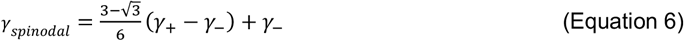

Thus, for a symmetric double-well function, it is not possible to set *γ*_*–*_, *γ*_*+*_, and *γ*_*spinodal*_ independently— two concentrations define the third (Figure B1C). This property is useful for determining whether a free energy function such as Equation 1 is appropriate for a phase-separated system.

**Figure. B1.**
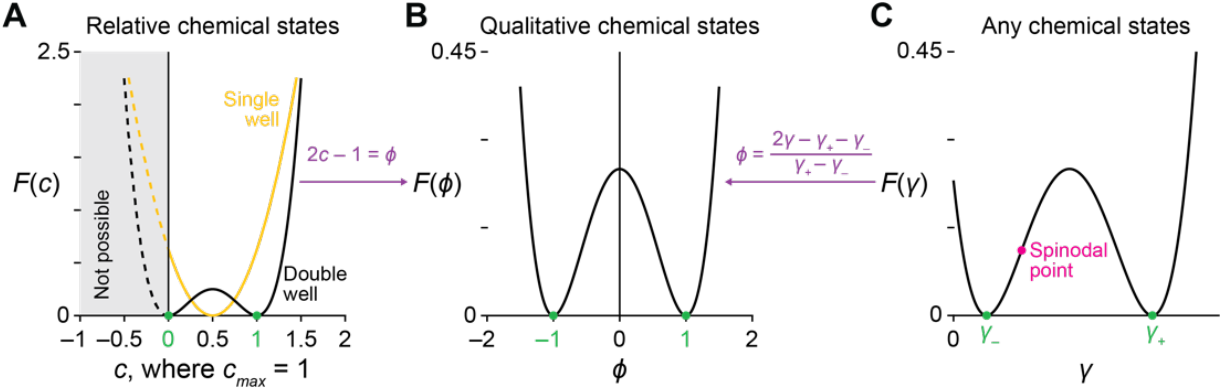
The Cahn-Hilliard double-well free energy function (*F*) accommodates differences in location and scale. (A) Comparison of a single-well function (yellow, with *c*_*avg*_ = 0.5) and a double-well function (black, stable states at 0 and 1 in green). (B) The double-well function of (A) is made symmetric about the y-axis with linear shifts and scaling (purple). (C) Any pair of chemical states (*γ*_*–*_ and *γ*_*+*_ in green) is similarly shifted and scaled to (B). The spinodal point (magenta) is constrained.

### Box 2. Dimensional analysis of the Cahn-Hilliard equation

For qualitative chemical states (*ϕ* = –1, 1 [=] dimensionless; Box 1), the Cahn-Hilliard equation is succinctly defined as:

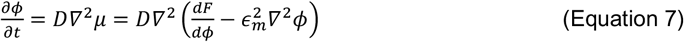

where *D* is diffusivity ([=] length^2^/time), *∇*^2^ is the Laplacian, *µ* is a generalized chemical potential ([=] energy nondimensionalized by *k*_*B*_*T*; Box 1), 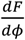 is the first derivative for Equation 4 ([=] nondimensionalized energy), and ϵ_*m*_ is the “interfacial energy coefficient”, a transition distance ([=] length•{nondimensionalized energy}^1/2^) operationally defined as:

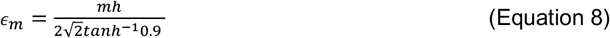

where *h* is the fractional mesh size of the spatial domain (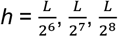, etc. where *L* is the size of the spatial domain [=] length) and *m* is the number of mesh points over which the interface exists ([=] dimensionless). (The 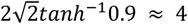 normalization arises from the equilibrium one-dimensional solution^27^ for an infinite interface, 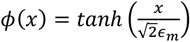, when considered over the range of *ϕ* = –0.9 to *ϕ* = 0.9. Thus, *ϵ*_4_ ≈ *h* and *ϵ*_8_ ≈ 2*h*.) In Equation 7, 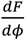 in *µ* captures departures of *F*(*ϕ*) from its thermodynamic minima (Figure B1B), and the 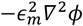 term generalizes *µ* to approximate interfacial surface energy for the diffuse interface between the two chemical states.^17^ Solving for an unknown *ϵ*_*m*_ is equivalent to defining *m* for a given *h*.

The generalized chemical potential *µ* is defined in terms of *ϕ* and normalized in such a way that its second-order changes occur on the same order of magnitude as changes in *ϕ* itself at all times (Box 1). Therefore, the effect of these field variables cancels and by scaling analysis:

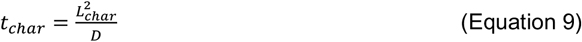

The solvers in this work set *D* = 1 for the general solution case. The actual physical dimensions of *D, ϵ*_*m*_, and *h* set different characteristic times (*t*_*char*_) depending on what length scale that diffusion is operating. For one-dimensional diffusion of *ϕ* across the operational interface in Equation 8:

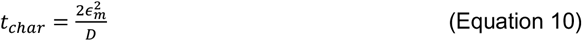

and for two-dimensional diffusion in the overall system:

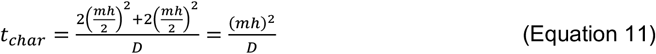

Taking a 3.2 µm × 3.2 µm patch of chromatin with *D* = 10^−3^ µm^2^/second,^28^ a 2^8^ = 256-element mesh, and *ϵ*_4_ ≈ *h* = 3.2 µ*m*/2^8^ = 0.0125 µ*m* yields *t*_*char*_ = 0.312 seconds for diffusion across the interface and *t*_*char*_ = 2.84 hours for diffusion across the system. (Note that for a same-sized patch of cytoplasm with *D* ~ 10 µm^2^/second,^29^ diffusion across the system would occur with *t*_*char*_ = 1.02 seconds.) Reciprocally, fixing a timeframe of interest defines a characteristic length scale: 10 minutes of chromatin diffusion will influence distances of biomolecular condensates on the order of 775 nm.

### Box 3. Nonlinear multigrid (NMG) and scalar auxiliary variable (SAV) approaches to solving the Cahn-Hilliard equation

In the NMG approach, spatial derivatives are cast as centered-difference approximations, and the method uses a semi-implicit time discretization with a nonlinear convex-splitting scheme for the free energy (Equations 1 and 4). This combined scheme ensures that a discrete version of the free energy is always decreasing and thus stable with time. NMG solves the nonlinear discrete equations at the implicit time level using the full approximation storage (FAS) multigrid method.^30^ The discrete system is solved iteratively on a hierarchy of square 2^*n*^ meshes (*n* = 6–9) that are obtained by successively coarsening the original mesh by factors of two in each direction. Smoothing is used to remove high frequencies and ensure accurate approximations of the full solution on the coarser meshes, as opposed to the error in the linear multigrid method. Coarse grid corrections are interpolated back to the fine grid and the corrected solution is smoothed. The nonlinearity is handled using a local linearization with Newton’s method and a pointwise Gauss–Seidel relaxation scheme is used as the smoother.^18,31^ This process is repeated recursively until convergence at each time step. NMG is more stable and efficient than standard Newton-type methods that rely on global linearizations (Figure 1) or algorithms that treat the nonlinear terms as forcing functions.^31^

**Figure 1.**
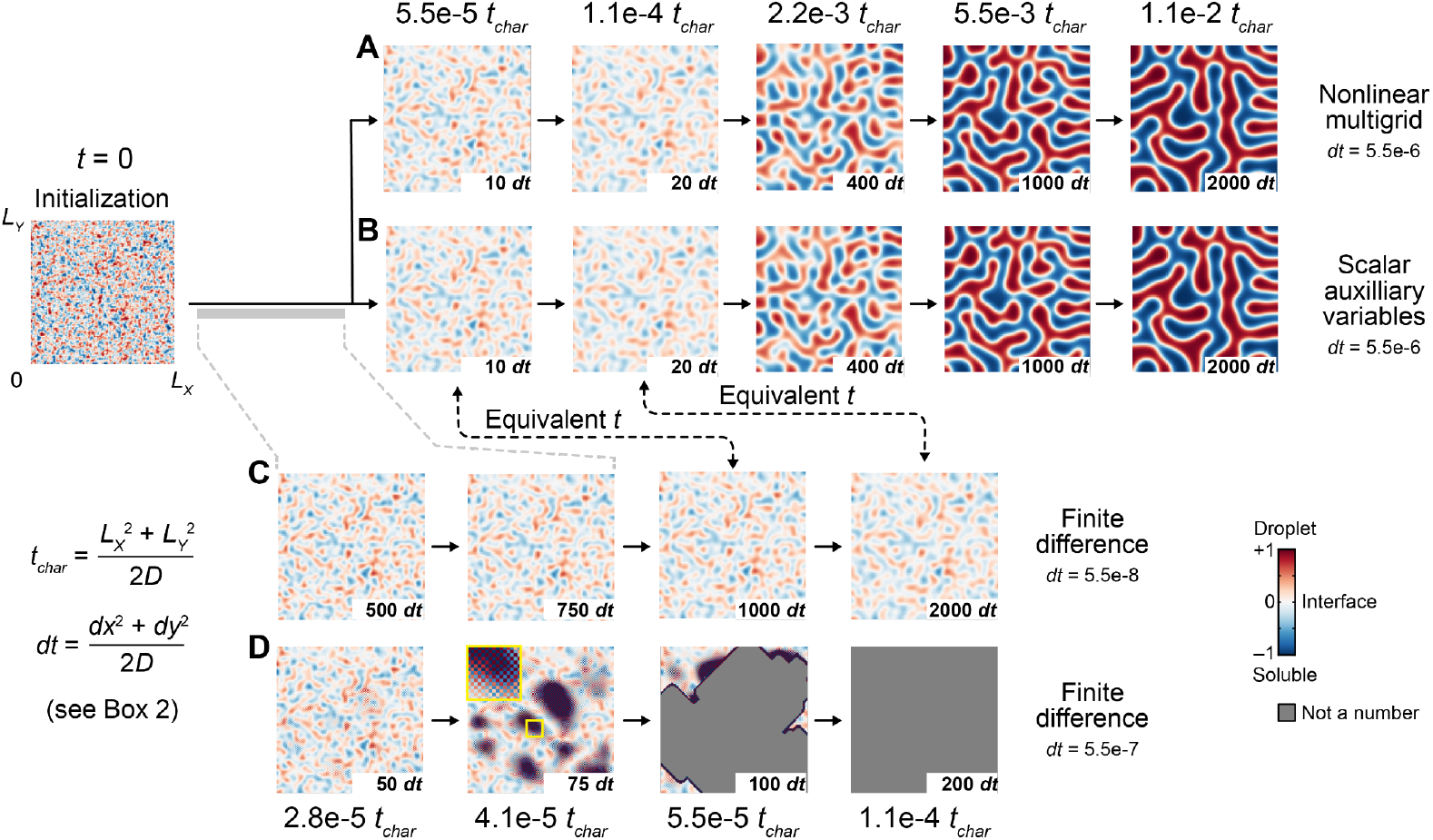
Practical simulations of the Cahn-Hilliard equation require customized solvers. (A and B) Time evolution of spinodal decomposition solved by nonlinear multigrid (A) or scalar auxiliary variables (B) over 1.1e-2 characteristic times (*t*_*char*_) defined by the length of X and Y spatial domains (*L*_*X*_, *L*_*Y*_) and system diffusivity (*D*). Both solvers used a 5.5e-6 time step (*dt*) defined by the mesh size in *X* and *Y* (*dx, dy*) and *D*. (C) Time evolution for the same spinodal decomposition solved by finite difference using a 100-fold smaller *dt* (5.5e-8) for 1% of the duration in (A) and (B). (D) Numerical instabilities triggered by the finite difference solver when using a 10-fold smaller *dt* (5.5e-7) than in (A) and (B). Defects appear as checkerboards in the mesh (inset) and become undefined (gray). Simulations were initialized as a random mixture of ±1 chemical states on a 2^7^ square mesh and smoothed before iterating with Neumann boundary conditions (STAR Methods). Alternative initializations and boundary conditions are shown in Figure S1. See also Figure S1.

For SAV, a new scalar variable is introduced in terms of the integrated non-gradient part of the free energy (*F*(*ϕ*) of Equation 7). Equation 7 is then appended with an additional ordinary differential equation for this scalar variable.^32^ Straightforward time discretizations of the reformulated system lead to decoupled linear, elliptic partial differential equations with constant coefficients. The decoupled equations are efficiently solved using the discrete Fourier transform^33^ for appropriate boundary conditions (periodic or Neumann). Like NMG, SAV is also energy stable in that the reformulated discrete energy, containing the scalar “auxiliary” variable, does not increase with time. To ensure that the reformulated discrete energy is consistent with the original system energy, a relaxation scheme is used to penalize discrepancies that may arise between the reformulated and original energies. For further details, see Ref. ^34^ and STAR Methods.

## RESULTS

### Modeling emergent subcellular compartments with the Cahn-Hilliard equation

Systems models of cell biology routinely encode nuclei, mitochondria, endosomes, and other organelles as subcompartments that are static and prespecified by membranes.^35–38^ Biomolecular condensates also compartmentalize but with shapes and sizes that are much more dynamic because of interfacial phenomena between condensed and soluble phases.^21,39^ Mathematically, interfacial free energy can be formally coupled to the energetics of weak intermolecular interactions^39^ (Box 1) and diffusion in space and time through the Cahn-Hilliard equation (Box 2).^17^ Dynamics of the Cahn-Hilliard equation naturally evolve to an equilibrium solution that minimizes the free energy of the overall system. The Cahn-Hilliard equation also conserves mass—solute diffuses freely within the system and can transition between condensed and soluble chemical states (Box 1), but the overall amount remains constant. This is advantageous for modeling biomolecular condensates that evolve on time scales of seconds to minutes (Box 2).^21,39^

### Numerical solvers for the Cahn-Hilliard equation

The energy-minimizing and mass-conserving properties of the Cahn-Hilliard equation set criteria for the accuracy of its numerical solutions. Computational challenges arise from the interfacial free energy term embedded within the generalized chemical potential (Equation 7), which results in a fourth-order spatial derivative that is problematic for standard solvers. The Cahn-Hilliard equation is also intrinsically stiff, because the fast dynamics of interfaces must be considered along with the slower dynamics of phases (Box 2). Maintaining energy dissipation and mass stability using standard methods requires very-fine spatial meshes, very-small time steps, or global linearizations, any of which adds computational burden.

Various custom methods solve the Cahn-Hilliard equation accurately and efficiently through different numerical schemes,^40^ but none are available with biology-focused users in mind. We thus developed generalized code for one finite-difference method and one spectral method in three widely used programming languages: Python,^41^ MATLAB, and Julia.^42^ Our finite-difference method adapts the nonlinear multigrid (NMG; Box 3) approach of Lee et al.^18^ The complementary spectral method uses the scalar auxiliary variable (SAV; Box 3) approach to gradient flows by Shen et al.,^32^ which can be similarly accurate and efficient. Together, NMG and SAV provide a versatile pair of Cahn-Hilliard solvers for biomolecular condensate systems.

Because NMG and SAV are very different algorithmically (Box 3), it was important to confirm that both packages yield the same numerical solutions. As an initial condition, we used a smoothed random mesh of ±1 chemical states (Figure 1, left, Box 1, and STAR Methods), which spontaneously undergoes a spinodal decomposition into two phases. After ~1% of characteristic time for the system (*t*_*char*_; Equation 11), both NMG and SAV converged to a slowly evolving solution with gradually decreasing energy and no overall change in mass (Figures 1A, 1B, S1A, and S1B). The resulting spatial pattern resembles the trajectory of biomolecular condensates triggered to self-assemble on lipid bilayers.^43^ We compared results of NMG and SAV to a standard finite-difference implementation of the forward Euler method and successfully captured the first 0.01% of *t*_*char*_ by taking 100-fold smaller time steps (*dt*; Figure 1C). Any larger time steps caused spatial defects in the finite-difference solution along with drastic deviations in energy and mass compared to the true solution (Figures 1D, S1A, and S1B). These results, plus others with alternate boundary conditions and smoothing initializations (Figures S1E–H), verified the self-consistency of our NMG–SAV implementations and reinforced the general need for them.

To compare run times between solvers and among computing environments, we re-simulated ~1% of *t*_*char*_ for three different spinodal initial conditions and two common boundary conditions: i) periodic, in which efflux from one edge is reflected as influx from the opposite edge, and ii) Neumann, or no flux along any edge (Figures 2A, S2A, and S2B). In Python, a language that many computational systems biologists consider numerically superior to alternatives (i.e., R), we found that NMG was largely unusable, requiring 10^5^–10^6^ seconds (by extrapolation) for each simulation (Figure 2B). The same implementation of NMG was 1000-fold faster in MATLAB and Julia for both boundary conditions. We attribute these differences to a non-tail recursive function in the NMG solver that is particularly suboptimal for Python.^18^ In contrast, SAV implementations yielded similar run times across the computing environments due to shared use of an efficient fast Fourier transform (FFT) algorithm for solving the elliptical equations posed by SAV (Box 3). Also, SAV performance depended on the boundary conditions—unlike NMG, Neumann simulations were handled less efficiently than spinodal decompositions that were periodic (Figure 2C; *P* = 5.3e-05). Due to the periodicity of the FFT, SAV solutions with Neumann boundary conditions must be solved by reflecting the spatial domain at the boundary, which increases the computations by fourfold (Figure 2C, inset). These overall trends persisted for mesh sizes ranging from 2^6^ × 2^6^ to 2^9^ × 2^9^ (Figures S2C and S2D). Because of their high initial spatial frequencies, spinodal decompositions are among the most difficult Cahn-Hilliard solutions, making them useful to discern whether the time step is appropriate for a given mesh size.

**Figure 2.**
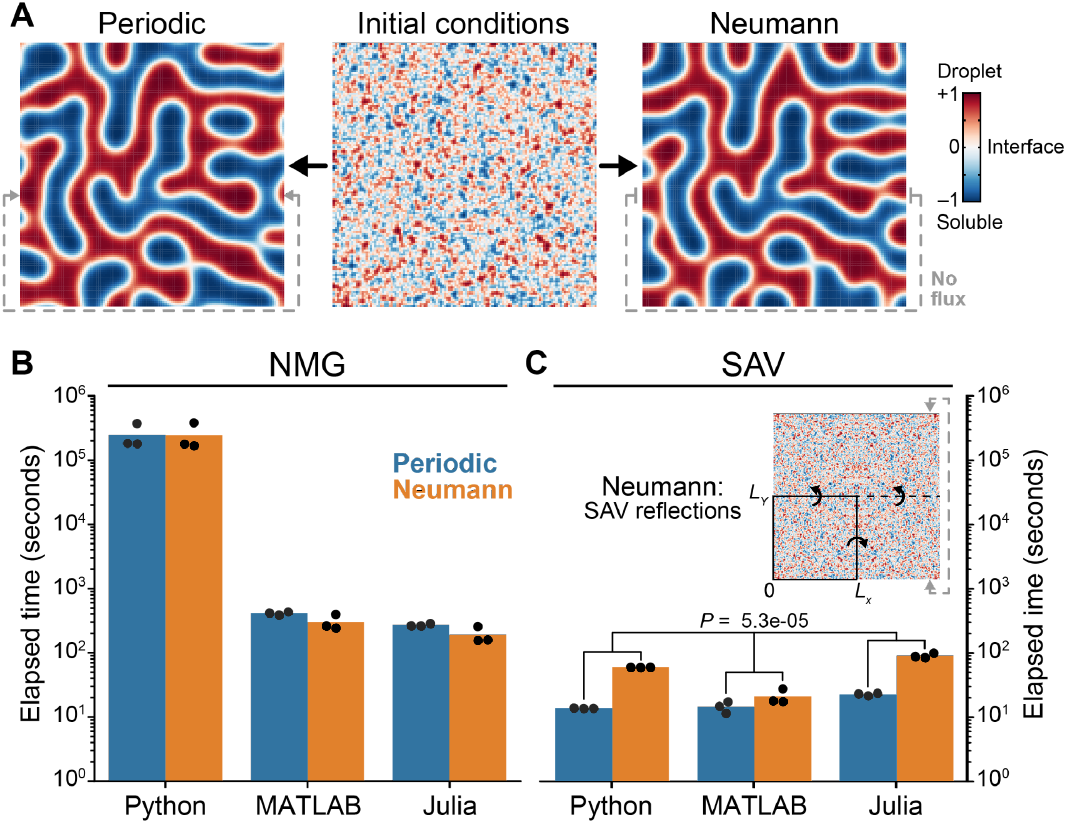
Computing performance depends on the programming language for NMG and the boundary condition for SAV. (A) Snapshots of spinodal decomposition when boundary conditions are periodic (left), whereby flux exiting one edge enters on the opposite edge, or Neumann (right) specifying zero flux at the edges. (B and C) Runtime performance for NMG (B) and SAV (C) solvers in Python, MATLAB, and Julia for spinodal decompositions initialized with *N* = 3 random mixtures of ±1 chemical states (25:75, 50:50, and 75:25) and either periodic or Neumann boundary conditions. For SAV, Neumann boundary conditions require *X* and *Y* reflections to re-establish periodic boundaries for the expanded spatial domain (C, inset). Boundary conditions in (C) were tested by multiway ANOVA with boundary condition and programming language as factors. All simulations were performed on a Linux x86_64 processor with 100 GB RAM on a 2^7^ × 2^7^ mesh (*L*_*X*_ = *L*_*Y*_ = 1) for 2000 time steps (*dt* = 5.5e-6) and *ϵ*_*m*=8_. See also Figure S2.

### Long-term Cahn-Hilliard simulations define universal parameters for condensate stability

The Cahn-Hilliard equation is handled non-dimensionally, but it formalizes concrete relationships between space, time, concentration, and diffusivity that are frameable in absolute terms (Boxes 1 and 2). The one unknown parameter is the interfacial energy coefficient defining the transition distance between phases (*ϵ*_*m*_; Equations 7 and 8), raising the practical question of how to constrain it for a specific biological system. We considered exploiting the criticality of Cahn-Hilliard fluids, whereby small phase-separated droplets dissolve into the bulk when there is not enough mass to retain them at a given surface tension (Figure 3A). For any *ϵ*_*m*_, there exists a critical droplet size below which droplets will not persist over time (File S1). The radius of this droplet may be defined in terms of its critical initial radius (*R*_*i*_) at *t* = 0 or its critical equilibrium radius (*R*_*eq*_) as *t* → ∞. Although there is theory linking *ϵ*_*m*_ to *R*_*i*_ under prescribed assumptions,^44^ the relationship between *ϵ*_*m*_ and *R*_*eq*_ is unknown but important because *R*_*eq*_ is estimable by experiments.

**Figure 3.**
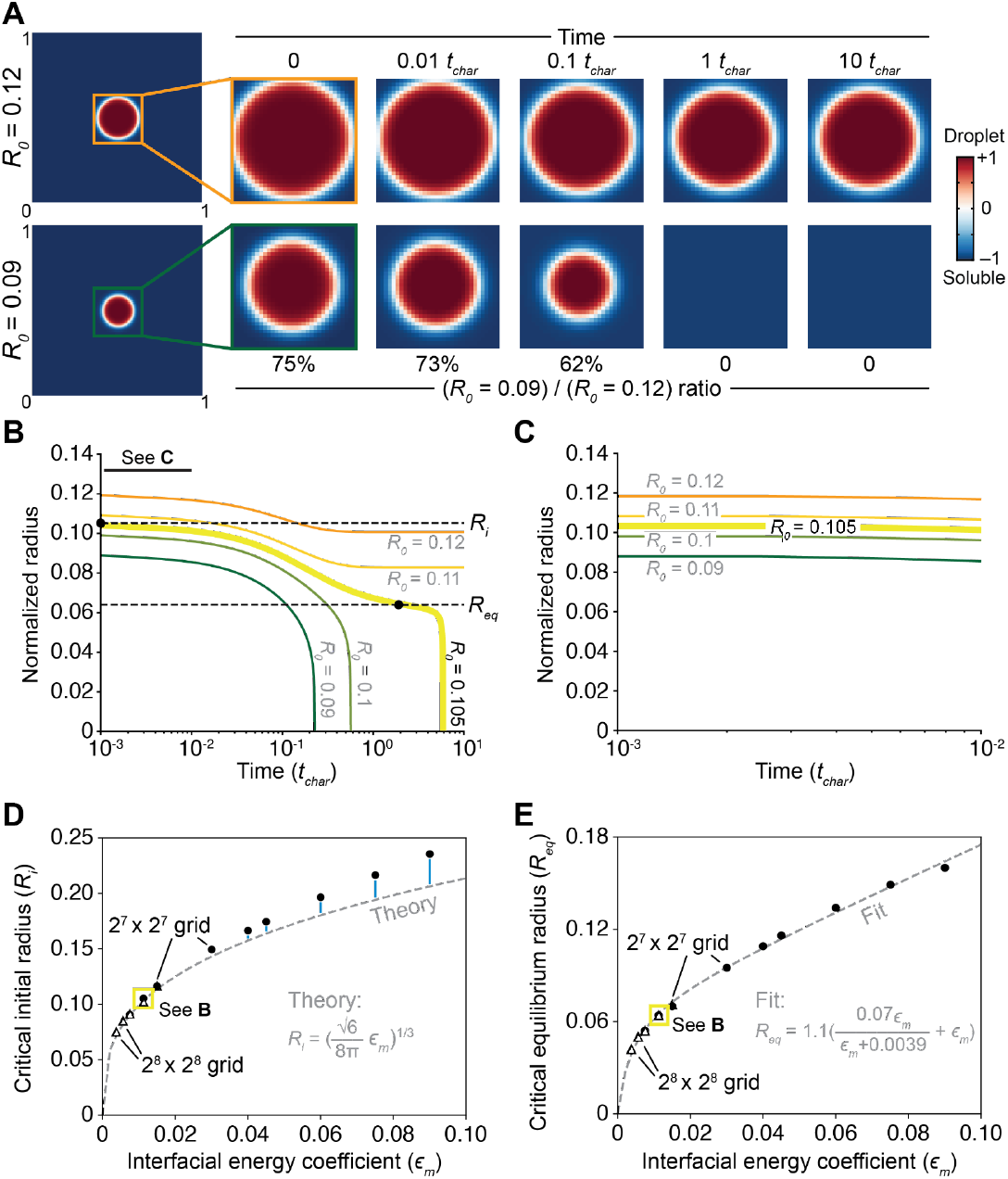
Multithreaded Cahn-Hilliard simulations define the relationship between *ϵ*_*m*_ and critical droplet radii. (A) Simulated dynamics for two normalized initial droplet radii (*R*_*0*_) on a 2^8^ × 2^8^ mesh (*L*_*X*_ = *L*_*Y*_ = 1) for 400,000 time steps (*dt* = 2.5e-5) and *ϵ*_*m*=12_ ≈ 0.011. The interface between the circular droplet and soluble phase was initialized to the hyperbolic tangent for an infinite interface (Box 2 and STAR Methods). The proportion of *R*_*0*_ = 0.09 relative to *R*_*0*_ = 0.12 is shown for each characteristic time point (*t*_*char*_). (B) Scan of *R*_*0*_ to determine the critical initial radius (*R*_*i*_) and the critical equilibrium radius (*R*_*eq*_) for a given *ϵ*_*m*_. *R*_*i*_ was calculated at *t* = 0 between the largest *R*_*0*_ that dissolves away (yellow) and the smallest *R*_*0*_ that persists (marigold). *R*_*eq*_ was estimated from the inflection point of the largest dissolving droplet (STAR Methods). (C) Enlargement of the first 0.1% of the time simulated in (C). (D and E) Calculation of *R*_*i*_ (D) and *R*_*eq*_ (E) as in (B) from *ϵ*_*m*=4_ on a 2^8^ × 2^8^ mesh (= 0.0037) to *ϵ*_*m*=48_ on a 2^7^ × 2^7^ mesh (= 0.0901). Calculations for *ϵ*_*m*=8,12,16_ on a 2^8^ × 2^8^ mesh (triangles) were repeated for *ϵ*_*m*=4,6,8_ on a 2^7^ × 2^7^ mesh (circles) to confirm overlap. In (D), calculations are compared to a prior theory^44^ (inset) relating *R*_*i*_ and *ϵ*_*m*_; systematic deviations are indicated in blue. In (E), calculations are compared to a hyperbolic-to-linear fit (inset); goodness-of-fit was 0.997. The approximate condition in (B) is highlighted (yellow box). See also Figure S3.

Defining critical radii requires long simulations, because droplets near the bifurcation often do not dissolve until one *t*_*char*_ or more has passed (Figure 3A). We systematically explored a range of starting non-dimensional radii (*R*_*0*_, which is normalized to the overall spatial domain) and simulated up to 10*t*_*char*_ with the stable NMG–SAV solvers by multithreading individual runs on a supercomputer. *R*_*i*_ was defined as the midpoint between the smallest *R*_*0*_ that did not dissolve and the largest *R*_*0*_ that did, whereas *R*_*eq*_ was approximated at the inflection point in time when the largest dissolving *R*_*0*_ starts to vanish (Figure 3B). Notably, none of these changes were evident with the first 1% of *t*_*char*_ (Figure 3C), reinforcing the need for numerically stable, efficient, and accurate solvers as provided here (Figure 1). Solutions for *R*_*i*_ and *R*_*eq*_ at a specific *ϵ*_*m*_ did not change if the starting radius was located off center in the spatial domain or split across the domain when boundary conditions were periodic (Figure S3A and S3B). Thus, *R*_*i*_ and *R*_*eq*_ are fundamentally linked to the *ϵ*_*m*_ of a Cahn-Hilliard system.

By repeating *R*_*0*_ sweeps across many *ϵ*_*m*_ values, we numerically defined the general relationship between *ϵ*_*m*_ and *R*_*i*_ or *R*_*eq*_ on a 2^7^ × 2^7^-element mesh (Figures 3D and 3E). For small *ϵ*_*m*_ values, 2^7^ elements were insufficient to specify the interfacial transition accurately (*m* ≥ 4 mesh points is recommended). Therefore, a 2^8^ × 2^8^-element mesh was substituted with enough overlap at intermediate *ϵ*_*m*_ values to confirm that results were identical. We observed excellent concordance between an approximation relating *R*_*i*_ and *ϵ*_*m*_ when *ϵ*_*m*_ values were small (Figure 3D), corroborating the underlying assumptions of this derivation.^44^ At large *ϵ*_*m*_ values, however, there was a systematic underestimate by the theory, consistent with the omission of higher-order terms in its derivation. For *R*_*eq*_, we sought an empirical regression that was simple and accurate, finding that a three-parameter hyperbolic-to-linear equation^45^ was best among alternatives (Figures 3E and S3C). Together, these relationships provide a reference for fully parameterizing any Cahn-Hilliard system given *R*_*i*_ or *R*_*eq*_.

### Cahn-Hilliard abstraction of the chromosomal passenger complex

We next sought a biological setting to apply the NMG–SAV solvers and Cahn-Hilliard constraints. In cells, biomolecular condensates form in the nucleus upon multivalent interactions with DNA.^10^ One highly dynamic example of multivalent binding involves the chromosomal passenger complex (CPC), a multi-protein assembly that is recruited to mitotic chromosomes between prophase and metaphase.^46^

CPC binding occurs through two chromatin modifications that overlap at the inner centromere between kinetochores of the sister chromatids (Figure 4A). One of the modifications (phospho-histone H3 Thr3, pH3T3) emanates from a kinase bound to cohesin, which holds together the sister-chromatid arms. The monovalent affinity^47^ of the pH3T3-CPC interaction is strong enough to recruit some CPC between sister chromatids in prophase, but only CPC recruited to the inner centromere, where there is a partially overlapping second histone phosphorylation, persists into metaphase.^48^

**Figure 4.**
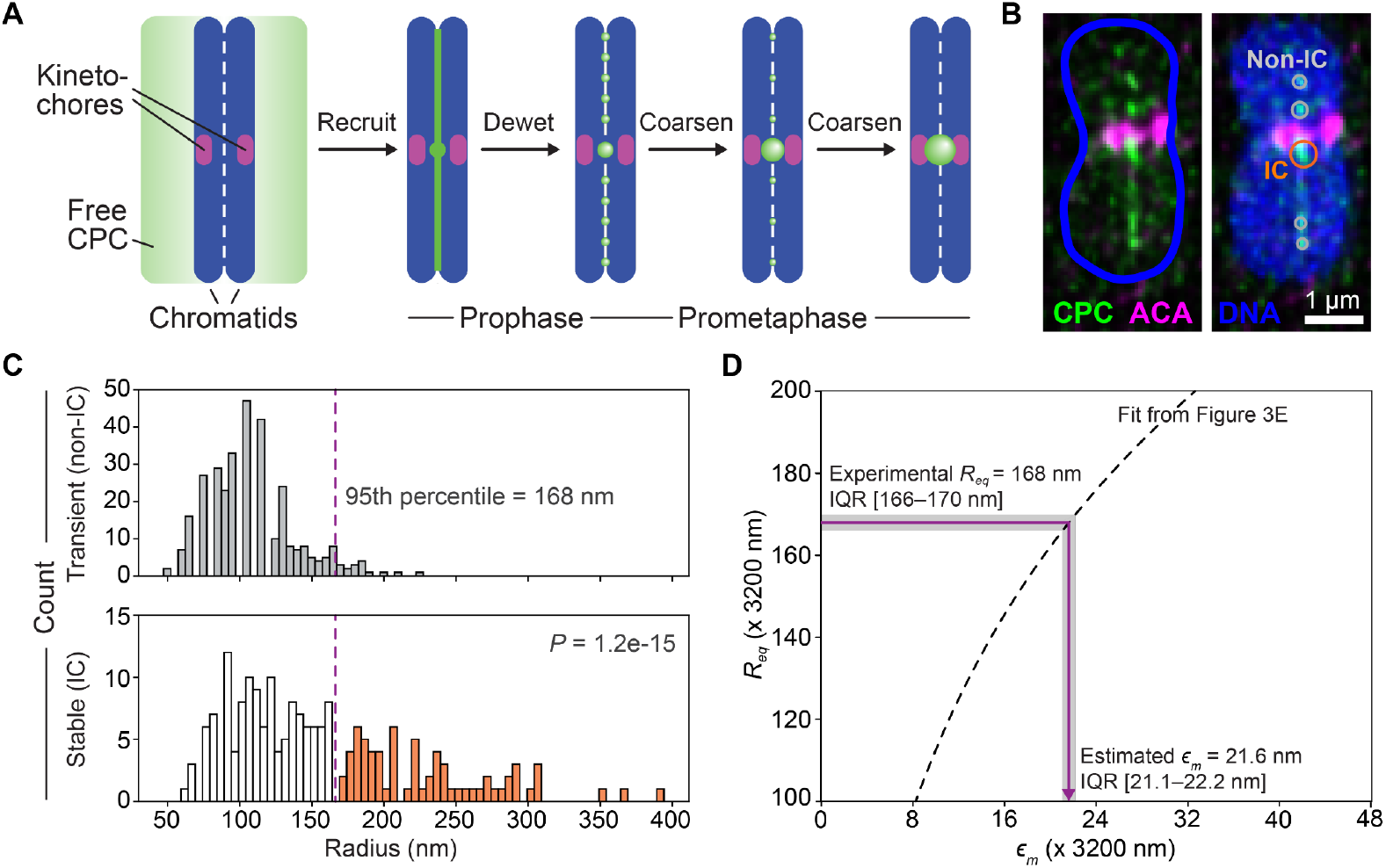
Chromatin-bound foci outside the inner centromere (IC) define a critical equilibrium radius (*R*_*eq*_) for the chromosomal passenger complex (CPC). (A) CPC recruitment to the IC during prophase and prometaphase. Freely diffusible CPC (green) and chromatin bound CPC is recruited strongly to the IC between kinetochores (magenta) and less so between sister chromatids. During late prophase, CPC dewets into discrete foci, which coarsen until all CPC is IC-localized by prometaphase. (B) Immunofluorescence illustration of CPC dewetting on a HeLa chromosome stained for AURKB (a CPC subunit; green), anti-centromeric antigen (ACA; magenta), and DAPI to label DNA (blue). ACA labeling of kinetochores distinguishes IC foci (orange) and non-IC foci (gray) of CPC during quantification (STAR Methods). (C) Histogram of non-IC foci (gray) and IC foci (white and orange) quantified by idealized circular radius from *N* = 221 chromosomes in 10 HeLa cells. The 95th percentile of the non-IC droplet size distribution defining *R*_*eq*_ is shown. The distribution of non-IC foci (*N* = 321) and IC foci (*N* = 195) were compared by KS test. (D) Estimation of *ϵ*_*m*_ from measured *R*_*eq*_ using the hyperbolic-to-linear regression (Figure 3E) scaled for a 3200-nm physical spatial domain. The interquartile range (IQR) was calculated by 50% subsampling of *N* = 10 HeLa cells for 100 iterations without replacement and propagating to the *ϵ*_*m*_ estimate. See also Figure S4.

Previously, Trivedi et al.^24^ reported that inner centromere-bound CPC reaches local concentrations high enough for its intrinsically disordered region (IDR) to form biomolecular condensates during mitosis. This claim was recently called into question^25^ based on new experiments involving how CPC foci form and dissolve at the inner centromere. We noted that the dynamics of the CPC from prophase to metaphase (idealized in Figure 4A) resembled two properties of a Cahn-Hilliard system. First, a Rayleigh-like instability^13,14^ breaks up monovalently bound CPC into smaller foci during prophase.

Then, an Ostwald-like ripening^15^ occurs during prometaphase, in which smaller CPC foci disappear as the larger inner centromere focus grows to become singular at the onset of metaphase. The µm length scale of changes (Figure 4B), the ~20-minute duration of (pro)metaphase,^49^ and the diffusivity of chromatin^28^ are all self-consistent within a Cahn-Hilliard framework (Box 2). Therefore, we modeled chromatin-bound CPC as a Cahn-Hilliard chemical state (+1; soluble CPC = –1) that could be tracked and tested by experiment.

We began with measurements in HeLa cells to build off earlier work^24,25,48^ and later examined the generality of the results using nontransformed cells of a different lineage. CPC binding was visualized on chromosomes by immunostaining mitotic spreads prepared after timed G2 release (STAR Methods). We stained chromosomes for AURKB (a subunit of the CPC) along with anti-centromeric antigen (ACA) to localize kinetochores and the inner centromere (Figure 4B). Most spreads showed CPC localized exclusively to the inner centromere (Figure 4A, right), but a fraction of cells exhibited multi-focal staining between sister chromatids (Figure 4B). The CPC characteristics of chromosomes in these multi-focal spreads were used to parameterize the Cahn-Hilliard equation and, eventually, to test it.

We constrained the interfacial energy coefficient by leveraging the general relationship between *ϵ*_*m*_ and *R*_*eq*_ (Figure 3E). We quantified the area of CPC foci from diffraction-limited confocal images that were deconvolved and z-projected (STAR Methods), converting the area to an equivalent radius for a circle (Figure 4B). Recognizing that CPC foci at the inner centromere persist and those elsewhere do not (Figure 4A), we stratified the measurements and noted that inner centromere foci had significantly larger radii (Figure 4C; *P* = 1.2e-15 by Kolmogorov–Smirnov (K-S) test). The partial size overlap with other foci was presumably because these inner centromere foci were still growing at the experimental endpoint (Figure 4A, middle), making it difficult for this distribution to define *R*_*eq*_. Therefore, we approached *R*_*eq*_ by instead using the distribution of non-inner centromere foci, all of which ultimately dissolve. Taking the upper 95th percentile of non-inner centromere foci, we estimated *R*_*eq*_ = 168 ± 2 nm for HeLa cells. We defined a square domain based on the ~95th percentile of measured chromosome arms (2 × 3.2 µm = 6.4 µm; Figure S4A) and used the relationship between *ϵ*_*m*_ and *R*_*eq*_ to define *ϵ*_*m*_ = 21.6 ± 0.6 nm (Figure 4D). Such a distance could be spanned by the ~32-nm single alpha-helical “dogleash” in between the chromatin-bound subunits of the CPC and kinase subunit, allowing the kinase to act inside or outside a putative condensate (Figure S4B).^50^

With a fully constrained system, we next examined the geometric requirements for a CPC “initial condition” at prophase (Figure 4A). Monovalent CPC recruitment was cast as a rectangle spanning the vertical domain with a prescribed width (*W*), upon which we overlaid recruitment to the inner centromere as a central circle with a radius of *R*_*IC*_ (Figure 5A). Initializing CPC as a rectangular crosshair^51^ quickly coarsened to a round droplet and yielded similar results to *R*_*IC*_ (File S2). We initialized Cahn-Hilliard simulations for large and small *R*_*IC*_ based on prior measurements of CPC foci at the inner centromere, and finer intervals of *W* were guided by CPC radii detected elsewhere (Figure 4C). Dynamics over the first 0.04*t*_*char*_ ≈ 7 minutes were considered realistic for prometaphase, but slowly evolving initial conditions were followed for nearly 70 minutes to define limiting behavior. The goal was to identify regimes of *R*_*IC*_ and *W* in which Rayleigh instability then Ostwald ripening was observable on the time scale of prometaphase.

**Figure 5.**
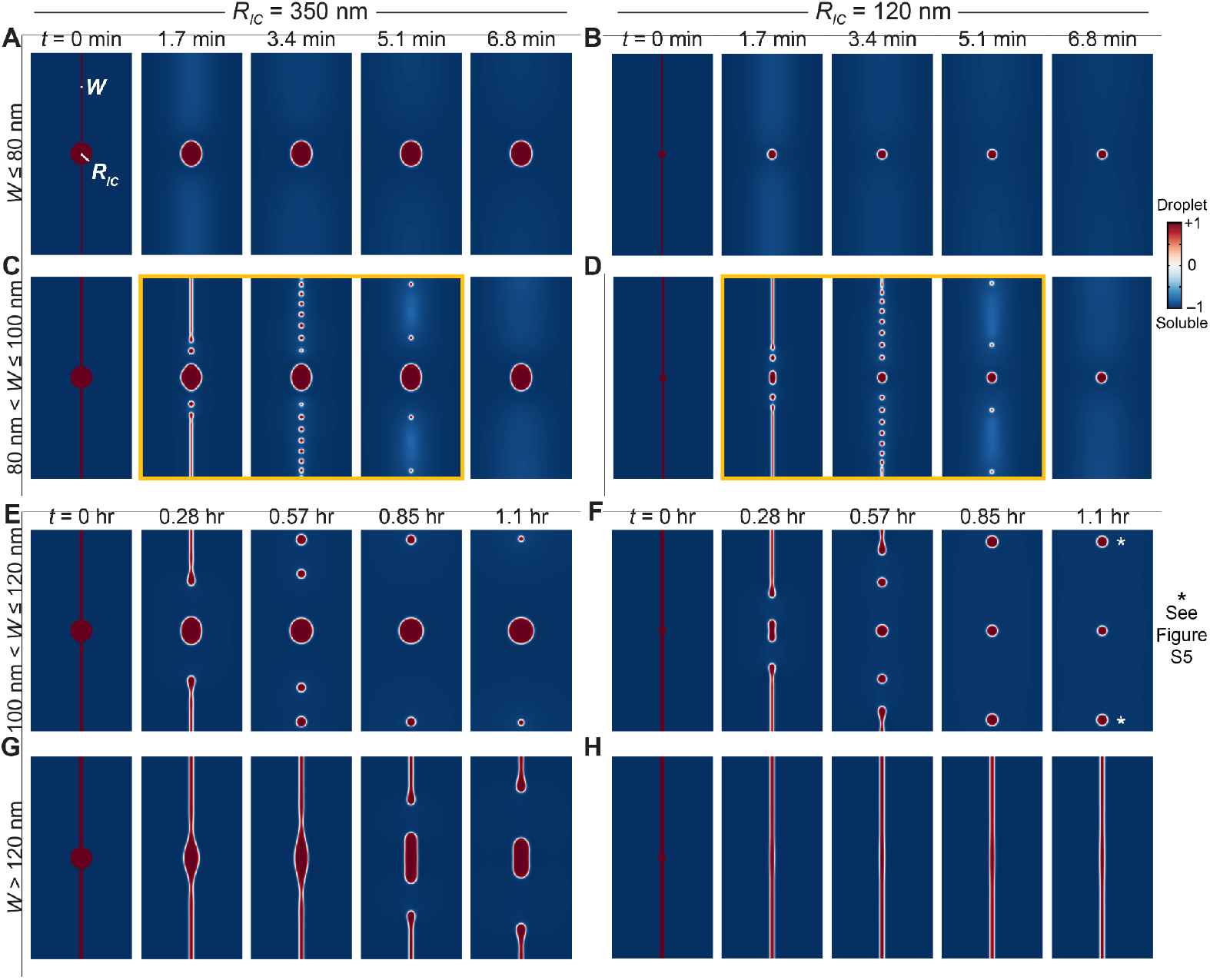
Cahn-Hilliard simulations predict extended multi-droplet regimes for physical values of CPC inner centromere radius (*R*_*IC*_) and pH3T3 width (*W*). (A–D) Dynamics at 1, 2, 3, and 4% *t*_*char*_ (corresponding to the indicated time in minutes based on chromatin diffusivity and mesh size; Equation 11) for *W* = 60 nm (A, B) or 90 nm (C, D) and *R*_*IC*_ = 350 nm (A, C) or 120 nm (B, D). Sustained droplets are highlighted in (C) and (D) (yellow). (E–H) Dynamics at 10, 20, 30, and 40% *t*_*char*_ for *W* = 120 nm (E, F) or 140 nm (G, H) and *R*_*IC*_ = 350 nm (E, G) or 120 nm (F, H). Sustained droplets (asterisks) are described further in Figure S5. Droplet patterns are representative for the intervals of *W* listed on the left. All simulations were performed on a Linux x86_64 processor with 200 GB RAM on a 2^9^ × 2^9^ mesh (*L*_*X*_ = *L*_*Y*_ = 2) for 26,214 (A–D) or 262,144 (E–H) time steps (*dt* = 1.53e-6). See also Figure S5.

Overall, we found that the emergent properties of the system depended disproportionately on *W*, with *R*_*IC*_ only affecting the final size of the central droplet except when *W* was exceedingly large (compare Figures 5A–5D to Figures 5E–5H). When *W* < 80 nm, axial droplets emerged and disappeared within 13 seconds, which would be too fast to see in fixed chromosomes (Figures 4B, 5A, and 5B). However, when *W* was increased to 80–100 nm, droplets persisted for several minutes before dissolving, providing enough time to capture them experimentally (Figures 4B, 5C, and 5D). This interval of *W* is notable because the cohesin holocomplex is 64 ± 7 nm long,^52^ and the pH3T3 kinase is tethered to cohesin by an extended linker on the N terminus that is likely unstructured.^53,54^ If this linker behaved as a random coil,^55^ its radius of gyration would be ~5 nm and thus add as much as ~20 nm to the effective width of cohesin for monovalent CPC recruitment, placing *W* in the 80–100 nm interval (Figure S5A).

Beyond 100 nm, the dynamics of the system were considerably slower. When *W* > 120 nm, the stripe was too stable and no droplets formed for over one hour (Figures 5G and 5H). Intriguingly, between 100 nm and 120 nm, a fourth regime slowly emerged, in which multiple droplets persisted for tens of minutes along the vertical axis (Figure 5E and 5F). This situation is not expected to arise ordinarily, but we identified one perturbation in the literature that may evoke it. Gascoigne et al.^56^ engineered HeLa cells to assemble nascent kinetochores ectopically throughout a chromosome, which should enlarge *W* by expanding the domain of inner centromere-like CPC recruitment (Figure S5B).

Revisiting the authors’ original images of CPC localization in metaphase-arrested cells, we identified lines of multiple foci on chromosomes with ectopic kinetochores, which were absent from controls (Figures S5C and S5D). Although not a formal prediction of the Cahn-Hilliard model, this observation built confidence in the physical dimensions of the system and our parameterization of *ϵ*_*m*_.

### Cahn-Hilliard models accurately predict spacing of CPC foci in multiple cellular contexts

We originally calibrated *ϵ*_*m*_ to the size distribution of bound CPC foci (Figures 4C and 4D). Since this calibration included no information about the location of foci along a HeLa chromosome, we reasoned that their spatial distribution would serve as an independent prediction of the resulting model. We focused on the multi-droplet regime based on prior structural arguments^52–55^ supporting feasibility (*W* = 80–100 nm; Figures 5C and 5D). In the idealized case, transient droplets appear and disappear with perfect symmetry on either side of the inner centromere (Figure 6A). Biologically, pH3T3 marks are spatially rougher^57^ and thus we displaced *W* by ±10 nm (one mesh point), which yielded an irregular pattern of longer-lived droplets (Figures 6B and File S3). Assuming that the immunofluorescence images were random snapshots of this dynamic, we built a gallery of Cahn-Hilliard movies for 10 *R*_*IC*_ values ranging from 50–350 nm (Figure 4C, bottom) and randomly sampled time points to build a predicted distribution of droplet-to-droplet distances (Figure 6B and STAR Methods). In parallel, we revisited the HeLa images and performed 300-nm-thick line scans of CPC–ACA staining between sister chromatids (Figure 6C) and found local maxima with a peak-finding algorithm (Figure 6D and STAR Methods). Using ACA to orient the inner centromere, we quantified peak-to-peak distances between CPC foci and estimated uncertainty in distance frequency by bootstrapping. The resulting histogram of measured peak-to-peak distances (Figure 6D) was directly comparable to the density of drop-to-drop distances predicted earlier by the Cahn-Hilliard model (Figure 6B).

**Figure 6.**
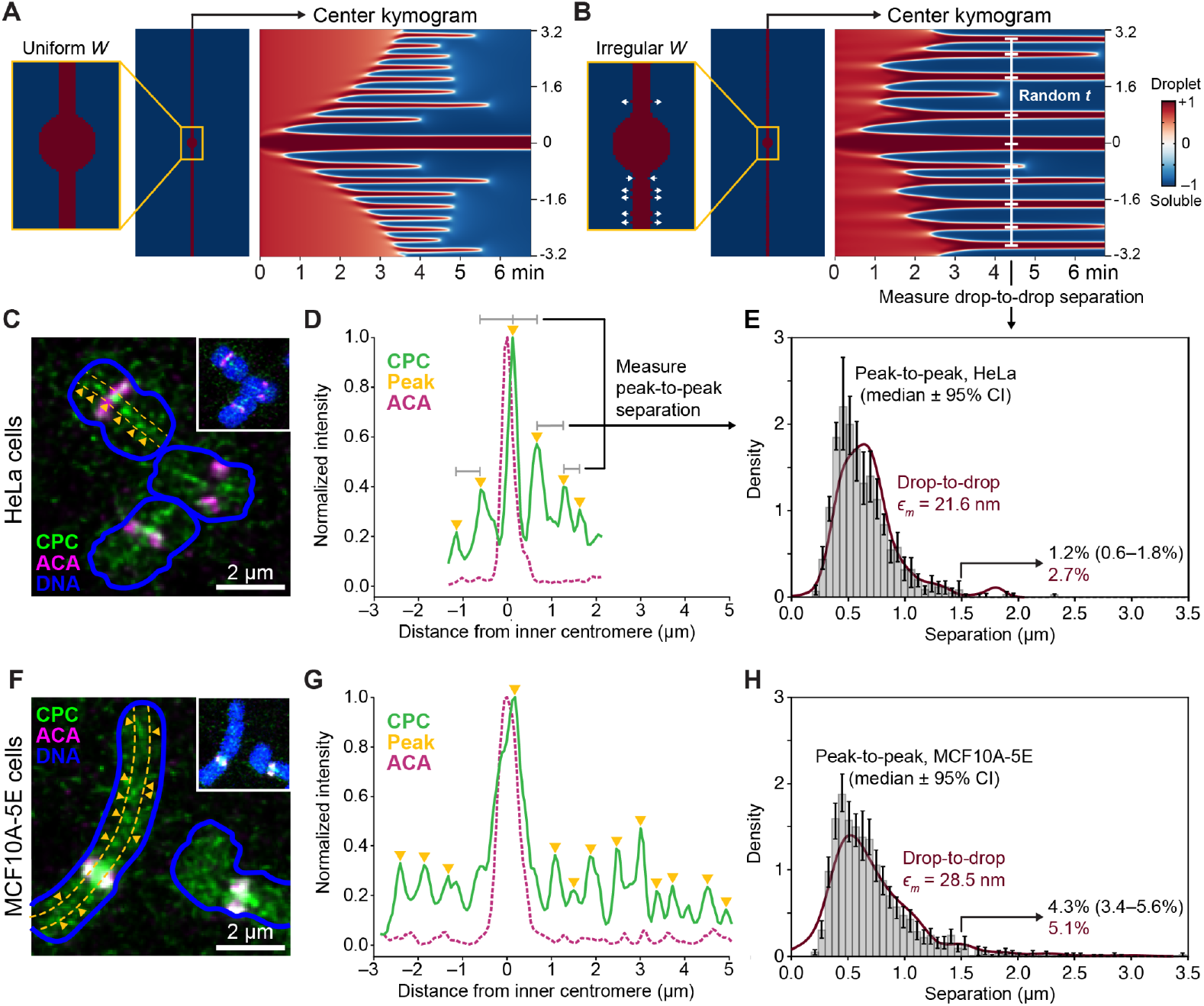
Chromatin-bound CPC behaves as a phase-separated Cahn-Hilliard fluid. (A and B) Kymogram illustration of central droplets for 6.8 minutes after initialization with *R*_*IC*_ = 150 nm and *W* = 90 nm (A) or 90 ± 10 nm (B, white arrows). Drop-to-drop distances were quantified at randomly selected times *t*. (C) Immunofluorescence staining of dewetted CPC on HeLa chromosomes after G2 release for 10 minutes. (D) Normalized intensity of AURKB and ACA for the HeLa chromosome in (C). (E) Comparison of bootstrapped peak-to-peak separation of CPC foci in HeLa cells (gray histogram) with drop-to-drop distances from Cahn-Hilliard simulations with *ϵ*_*m*_ = 21.6 nm (red line). (F) Immunofluorescence staining of dewetted CPC on MCF10A-5E chromosomes after G2 release for 70 minutes. (G) Normalized intensity of AURKB and ACA for the MCF10A-5E chromosome in (G). (H) Comparison of bootstrapped peak-to-peak separation of CPC foci in MCF10A-5E cells (gray histogram) with drop-to-drop distances from Cahn-Hilliard simulations with *ϵ*_*m*_ = 28.5nm (red line). For (C) and (F), cells were stained for AURKB (a CPC subunit; green), anti-centromeric antigen (ACA; magenta), and DAPI to label DNA (blue). A 300-nm thick axial line scan (dotted yellow line) was used to identify peaks of AURKB foci (yellow triangles; STAR Methods). For (D) and (G), normalized fluorescence intensities are shown relative to distance from the inner centromere based on the peak ACA location centered at zero. Peaks of AURKB foci (yellow triangles) were identified with a computational algorithm (STAR Methods) and quantified. For (E) and (H), data are shown as the median ± 95% bootstrapped confidence interval (CI) from *N* = 221 chromosomes in 10 cells (E) or 216 chromosomes in 50 cells (H). The fraction of predicted (red) and measured (black) separations greater than 1.5 µm is indicated. See also Figure S6.

Without any adjustments, we found that Cahn-Hilliard predictions of spatial frequency were nearly superimposable with CPC measurements in HeLa cells (Figure 6E). Separation distances of 500–600 nm were most common, and the right-tailed distribution decreased sharply with distance such that spacings were rare above 1 µm and almost nonexistent above 1.5 µm. By physically constraining a diffuse interface (*ϵ*_*m*_; Figures 4C and 4D) and an initial geometry (*W* and *R*_*IC*_; Figures 5C and 5D), the Cahn-Hilliard equation yields dynamics (Figure 6B) and length scales (Figure 6E) that are fully consistent with CPC biology.

Is *ϵ*_*m*_ a universal or context-specific property of the CPC? To address this question, we pivoted from HeLa cells, a highly aneuploid and divergent^58^ cervical cancer line, to the 5E subclone^59^ of MCF10A cells—a pseudo-diploid, non-cancerous^60^ breast epithelial line. Compared to HeLa, MCF10A-5E cells required much more time after G2 release to enter mitosis (STAR Methods). Further, CPC foci were most evident earlier in prophase, when chromosomes were not as fully condensed and thus more elongated (Figures 6F and S4A; *P* = 7.7e-48). We repeated the *R*_*eq*_ estimation for CPC as before and found that *ϵ*_*m*_ for MCF10A-5E cells was ~30% larger than for HeLa cells (28.5 nm ± 1.1 nm; Figures S6A and S6B). The increase may relate to overall chromatin compaction or differences in CPC modification state (see Discussion), but regardless the MCF10A-5E data indicate that the *ϵ*_*m*_ for CPC is not fixed.

The larger *ϵ*_*m*_ of MCF10A-5E simulations did not qualitatively impact whether multiple droplets formed over the same range of initializations (*W* = 80–100 nm; *R*_*IC*_ = 50–350 nm), but it did alter the droplet spacing. For the same predicted median spacing of ~640 nm, the predicted variance for MCF10A-5E droplets was greater, especially on the right tail of the distribution (Figure S6C). We repeated the line-scan and peak calls for CPC foci on MCF10A-5E chromosomes and observed the same exaggerated right tail compared to HeLa (Figures 6F, 6G, and S6D). When Cahn-Hilliard predictions and MCF10A-5E measurements were directly compared, there was excellent agreement, including more-frequent events in the 1–1.5 µm range and instances that extended beyond 1.5 µm (Figure 6H). We conclude that the CPC behaves as a Cahn-Hilliard fluid early in mitosis. More generally, the numerical methods gathered here enable a complementary validation strategy^25^ for biomolecular condensates that is purely mathematical.

## DISCUSSION

The Cahn-Hilliard equation emerged from chemical physics but has found application in many areas of biology.^1–6,8,9,17^ The refined NMG and SAV solvers presented in this work bring Cahn-Hilliard modeling closer to the realm of systems biology. The two numerical methods are not redundant but rather complementary. NMG is a more-established approach^31^ that uses its own defined functions recursively to solve the Cahn-Hilliard equation within a specified error tolerance. By checking against the Cahn-Hilliard equation at each time step, NMG acts as a standalone solver. The newer SAV approach^32^ changes the problem into a set of elliptical equations that are efficiently solved by a separate algorithm^33^ for discrete Fourier transforms. SAV is generally faster than NMG and represents the only practical option for Python. However, care must be taken to ensure that SAV stays true to the actual Cahn-Hilliard solution. Our SAV implementation incorporates relaxation^34^ to help with problematic initial conditions, like spinodal decomposition, and solution drift with very large time steps. When SAV performance is uncertain, NMG serves as a useful crosscheck for self-consistency.

We used Cahn-Hilliard simulations as a testbed for dewetting–coarsening of the CPC and found that its behavior was highly supportive of phase separation with *ϵ*_*m*_ specifying the diffuse interface width. By operating in physical units, parameter estimates were directly relatable to molecular entities. The linker subunit of the CPC has a ~32-nm long alpha helix^50^ preceded by an intrinsically disordered region that adds 5–14 nm depending on its phosphorylation state.^61^ Interestingly, neither of these regions is necessary for the CPC to form a condensate,^24^ raising the possibility that they may instead contribute to the magnitude of *ϵ*_*m*_. Although we did not detect cell-specific differences in overall phosphorylation of the CPC linker, another subunit was differentially altered, and there was a consistent 2–3-fold difference in overall abundance that might alternatively impact *ϵ*_*m*_ indirectly (Figures S6E and S6F). A long-term interest is to extend this framework to other phase-separated systems warranting compartmental models.^62,63^

Condensation of the CPC may be evolutionarily advantageous for the robustness it endows. In a Cahn-Hilliard system, larger droplets fuse with smaller droplets (or pull mass from them) until one droplet or zero droplets remains. The phenomenon acts as an analog-to-digital converter, which filters smaller, off-target droplets outside of the inner centromere. From this perspective, the system may employ two histone-binding activities emanating from orthogonal axes to ensure that the inner centromere generates a large enough *R*_*IC*_ so that it reliably dominates over alternatives.

A limitation of the standard Cahn-Hilliard equation (and, by extension, the NMG–SAV solvers) is the lack of source and sink terms for synthesis and degradation over time. Source–sink terms can be appended to the governing equation along with feedback,^3^ but given the number of possible configurations we opted for simplicity in the software packages. Constant mass is a reasonable assumption for the ~6-min dynamics of the CPC, although we recognize it may be less satisfactory for systems that evolve more slowly with time.

Biomolecular condensates are integrated or involved in many steps of cellular regulation. The properties of isolated condensates are well documented,^10–12^ but condensation is also a mechanism for conferring switch-like behavior to signaling pathways.^64,65^ Stitching Cahn-Hilliard dynamics into biochemical reaction-diffusion networks with these modern solvers is a next frontier for systems-biology modeling.

## RESOURCE AVAILABILITY

### Lead Contact

Further information and requests for resources and reagents should be directed to and will be fulfilled by the Lead Contact, Kevin A. Janes.

### Materials Availability

This study did not generate new unique reagents.

### Data and Code Availability

Code used to generate the results and figures in this paper is available on GitHub (https://github.com/JanesLab/GrovesSM_CahnHilliard).

Python, Julia, and MATLAB packages for modeling the Cahn-Hilliard equation, complete with tutorials, is available on GitHub (https://github.com/JanesLab/CahnHilliard_NMGSAV).

## ACKNOWLEDGMENTS

We thank Matthew Lazzara for critically reviewing the mathematical preliminaries of the manuscript and Iain Cheeseman for providing the original images of kinetochore-perturbed cells from Gascoigne et al.^56^ This work was initiated by a collaborative research supplement to the SASCO Cancer Systems Biology Research Center (3-U54-CA274499-02S1, K.A.J. and J.S.L.) and further supported by the National Cancer Institute (U54-CA274499, K.A.J.; P01-CA288662, J.S.L.), the University of Virginia Systems & Biomolecular Data Science training grant (T32-GM145443, K.A.J. and A.C.A.-Y.), the National Science Foundation (DMS-2309800, DMS-1763272, J.S.L.) and an ASPIRE Grant from the Susan G. Komen foundation (ASP241255843, K.A.J. and A.C.A.-Y.).

## AUTHOR CONTRIBUTIONS

Conceptualization: S.M.G., M.J.L., P.T.S., J.S.L., K.A.J. Data curation: S.M.G., K.A.J.

Formal analysis: S.M.G., M.J.L., J.S.L., K.A.J.

Funding acquisition: A.C.A.-Y., P.T.S., J.S.L., K.A.J.

Investigation: A.C.A.-Y., M.G.-R., K.A.J.

Methodology: S.M.G., M.J.L., J.S.L., K.A.J.

Project administration: J.S.L., K.A.J. Resources: P.T.S.

Software: S.M.G., M.J.L., K.A.J.

Supervision: J.S.L., K.A.J. Validation: A.C.A.-Y., M.G.-R., K.A.J.

Visualization: S.M.G., K.A.J.

Writing – original draft: S.M.G., M.J.L., J.S.L., K.A.J.

Writing – review & editing: S.M.G., M.J.L., A.C.A.-Y., M.G.-R., P.T.S., J.S.L., K.A.J.

## STAR METHODS

### EXPERIMENTAL MODEL AND STUDY PARTICIPANT DETAILS

#### Cell lines

HeLa cells (female) were cultured in Eagle’s Minimum Essential Medium (ATCC) supplemented with 10% fetal bovine serum (HyClone). MCF10A-5E cells (female) ^59^ were grown in Dulbecco’s modified Eagle’s medium/F-12 (Gibco) supplemented with 100 ng/ml cholera toxin (Sigma), 20 ng/ml epidermal growth factor (Peprotech), 10 mg/ml insulin (Sigma), 500 ng/ml hydrocortisone (Sigma) and 5% horse serum (Gibco). All base media were supplemented with 1% penicillin/streptomycin (Gibco) and all cultures were grown with 5% CO_2_ in a humidified incubator at 37ºC. By STR profiling (ATCC), HeLa cells were authenticated to be derived from RRID:CVCL_0030 (93% match), and MCF10A-5E cells were authenticated to be derived from RRID:CVCL_0598 (100% match).

## METHOD DETAILS

### NMG solver

We refactored the NMG algorithm of Lee et al.^18^ by hierarchically organizing local and global functions, integrating global functions with the SAV solver, and clarifying variable nomenclature usage. ‘CahnHilliard_NMG’ takes an initial chemical state field (*ϕ*_0_; Box 1) provided by the user or created by ‘ch_initialization’ and uses ‘nmg_solver’ with recursive calls to ‘nmg_vcycle’ that solve for *ϕ* at each time step *dt*. ‘CahnHilliard_NMG’ handles the following simulation parameters with the variable name and default value in parentheses: number of time iterations (t_iter = 1e3), the size of the time step (dt = 2.5e-5), the number of solver iterations per time step (solver_iter = 1e4), the absolute solver tolerance per time step (tol = 1e-5), the number of mesh points *m* for *ϵ*_*m*_ (m = 8), the boundary conditions (boundary = ‘periodic’), the number of smoothing relaxations done at the start and end of each coarsening–interpolation cycle (c_relax = 2), the domain size in terms of right and left coordinates of x and y (domain = [1 0 1 0]; note that [2 0 2 0] was used for the CPC simulations in Figures 4–6 and S4– S6), logicals to print residuals (printres = false) or *ϕ* at each time step (printphi = false), and the path name where *ϕ* will be output as a CSV (pathname = ‘cd’).

### Relaxed SAV algorithm and solver

For clarity, we retain the nomenclature for the relaxed SAV method of Jiang et al.^34^ The nonlinear free energy:

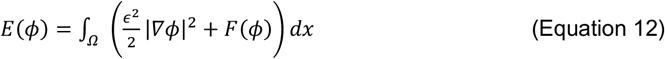

Is recast with the following SAV over the spatial domain *Ω* at time *t*:

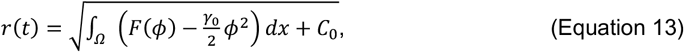

where 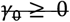 and *C*_*0*_ ≥ 0 ensure that the square root in *r*(*t*) is well defined. *γ*_0_ and *C*_*0*_ do not alter the underlying PDE but improve the unconditional energy stability of the numerical scheme. In the SAV code, we set *γ*_*0*_ = 2 and *C*_*0*_ = 1 as default values. Our empirical tests indicate these choices provide a good balance between accuracy and stability, which is consistent with previous results.^32,34,66^ We follow the regularized SAV framework of Chen et al. ^67^ and its relaxed extension from Jiang et al.^34^ by adding a quadratic term 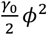 to the free energy *E*(*ϕ*), which is then subtracted after squaring the auxiliary variable *r*(*t*):

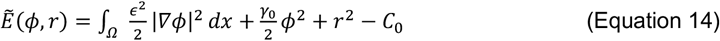

The auxiliary variable *r*(*t*) in Equation 13 together with the modified energy 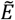 in Equation 14 yield a linear system of equations with constant coefficients when using Crank-Nicolson time discretization to solve for *ϕ* and *r*:

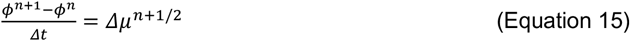

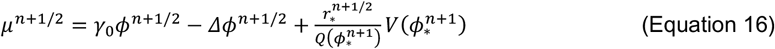

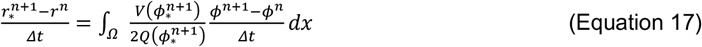

(Equation 17) where 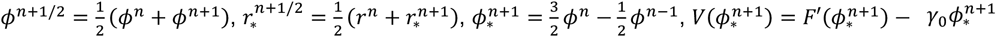, and 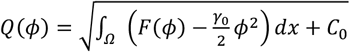. The terms 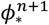 and 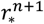 refer to provisional solutions for the next time step: *ϕ*^*n*+1^ and *r*^*n*+1^. The algorithm then applies a relaxation step to the provisional 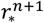:

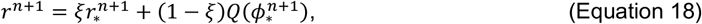

where *ξ* ∈ [0, 1] is the real root of the quadratic:

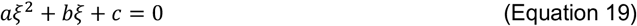

with:

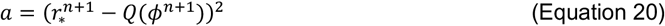

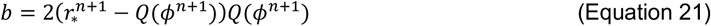

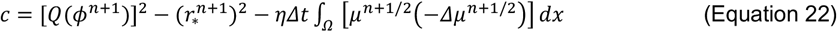

The term *η* ∈ [0, 1] scales the dissipation term from no dissipation (*η* = 0) to maximum dissipation (*η* = 1). In the SAV code, we take *η* = 0.95 as a default value, which was found to result in good accuracy and numerical stability.^34^ With Equations 18–22, *ξ* is computed to minimize |*r*^*n*+1^ − *Q*(*ϕ*^*n*+1^)| by solving the following optimization problem:

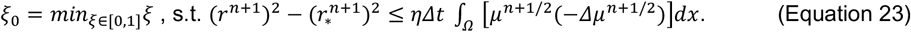

Once *r*^*n*+1^ is obtained through the above relaxation, *ϕ*^*n*+1^ is updated by Equation 15.

The SAV algorithm was coded to share global functions with the NMG solver as much as possible. ‘CahnHilliard_SAV’ takes an initial chemical state field (*ϕ*_*0*_; Box 1) provided by the user or created by ‘ch_initialization’ and uses ‘sav_solver’ [together with numpy.fft.fft2 (Python), fft2 (MATLAB), or fft (Julia FFTW) and other auxiliary functions in the package] to solve for *ϕ* at each time step *dt*. ‘CahnHilliard_SAV’ handles the following simulation parameters with the variable name and default value in parentheses: number of time iterations (t_iter = 1e3), the size of the time step (dt = 2.5e-5), the number of mesh points *m* for *ϵ*_*m*_ (m = 8), the boundary conditions (boundary = ‘periodic’), the domain size in terms of right and left coordinates of x and y (domain = [1 0 1 0]), logicals to print *ϕ* at each time step (printphi = false), the path name where *ϕ* will be output as a CSV (pathname = ‘cd’), the regularization parameter *C*_*0*_ (C0 = 1), the stabilization parameter *γ*_*0*_ (gamma0 = 2), the relaxation parameter *η* (eta = 0.95), and a logical for invoking relaxation (xi_flag = true).

### Finite difference solver

Outside of the Python, MATLAB, and Julia packages, we encoded a separate (not-recommended) set of functions in MATLAB that attempts to solve the Cahn-Hilliard equation by the forward Euler method of finite difference. ‘CahnHilliard_FD’ takes an initial chemical state field (*ϕ*_*0*_; Box 1) provided by the user or created by ‘ch_initialization’ and uses ‘fd_solver’ to estimate *ϕ*(*t*+*dt*) by linear extrapolation from *ϕ*(*t*). ‘CahnHilliard_FD’ handles the following simulation parameters with the variable name and default value in parentheses: number of time iterations (t_iter = 1e3), the size of the time step (dt = 2.5e-5), the spacing of timesteps that are printed or saved (dt_out = 10), the number of mesh points *m* for *ϵ*_*m*_ (m = 8), the boundary conditions (boundary = ‘periodic’), the number of smoothing relaxations done at the start of the simulation (c_relax = 2) and a logical variable to determine if the initial conditions are smoothed (presmooth = false), the domain size in terms of right and left coordinates of *x* and *y* (domain = [1 0 1 0]), logicals to print residuals (printres = false) or *ϕ* at each time step (printphi = false), and the path name where *ϕ* will be output as a CSV (pathname = ‘cd’).

### Spinodal decomposition

Square meshes of size 2^6^, 2^7^, 2^8^, or 2^9^ were randomly initialized with +1 at a probability of 25%, 50%, or 75% in Julia by randomizing +1 and –1 using the shuffle! function (random seed = 1234). The ‘relax’ function in the Julia package was then used to smooth the initial conditions with the following parameters: mu = zeros(GridSize,GridSize), dt = 6.25e-6, n_relax = 4, Lx = Ly = 1, m = 8, and boundary = “neumann”. These new smoothened initial conditions were used as *ϕ*_0_ for simulations. NMG used default values for tolerance and solver_iter. SAV used default values for *C*_*0*_, *γ*_*0*_, *η*, and regularization.

### Critical droplet simulations

Critical droplets were initialized on 2^7^ or 2^8^ square meshes with the ‘initialization’ function in the Julia package and method = “droplet”, which places a +1 circle with radius *R*_*0*_ in the center of the domain surrounded by –1 and the equilibrium interface 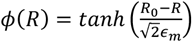 (Box 2). *ϵ*_*m*_ was altered by varying *m* = 4–48 and changing the mesh size (Equation 8). Simulations were performed with the ‘CahnHilliard_NMG’ function in the Julia package and tol = 1e-6, dt = 2.5e-5, solver_iter = 1e4 for 10 *t*_*char*_. Initial guesses of *R*_*0*_ were refined to three significant digits upon identifying the threshold of critical droplet behavior for a given *ϵ*_*m*_. Consistency was confirmed by i) repeating a subset of simulations with the ‘CahnHilliard_SAV’ function in the MATLAB package and ii) altering the position of the droplet (Figures S3A and S3B) with boundary = “periodic.”

### Immunofluorescence of chromosome spreads after G2 release

10-cm dishes of cells at 70% confluency were treated with 2 mM thymidine (Sigma) in fresh medium for 24 hours. Cells were washed three times with 5 ml PBS, and 9 µM RO-3306 (Selleckchem) was added with fresh media for an additional 24 hours. For HeLa, cells were again washed three times with 5 ml PBS and trypsinized. For MCF10A-5E, cells were trypsinized in the presence of 9 µM RO-3306, and pelleted cells were washed three times with 5 ml PBS. Pelleted cells were then transferred to fresh medium and incubated at 37°C for 10 minutes (HeLa) or 70 minutes (MCF10A-5E). Cells were again pelleted and then hypotonically swelled in 75 mM KCl, 0.8% Na Citrate, and H_2_O in a 1:1:1 ratio for 15 minutes at room temperature. Swollen cells were centrifuged onto coverslips at 67 rcf for 5 minutes with a Cytospin 4 (Thermo Shandon) and transferred to 6-well plates for staining. Samples were fixed with 2% paraformaldehyde (Thermo Fisher) for 20 minutes, blocked with 1% (w/v) bovine serum albumin (Fisher BioReagents) in PBS for 1 hour at room temperature, and stained for AURKB (BD Biosciences; 1:250 dilution), Centromere Antigen (Antibodies Incorporated, 1:500 dilution), and phospho-Histone H3 (Thr3) (Abcam; 1:250 dilution) in PBS + 0.1% Tween-20 (Sigma) (PBS-T) overnight at 4°C. Samples were washed three times with PBS-T for 5 minutes and then incubated for 1 hour at room temperature with the following secondary antibodies diluted in PBS-T: Alexa Fluor 488-conjugated goat anti-mouse (Thermo Fisher; 1:1000 dilution), Alexa Fluor 568-conjugated goat anti-human (Thermo Fisher; 1:1000 dilution), and Alexa Fluor 647-conjugated goat anti-rabbit (Thermo Fisher; 1:1000 dilution). Samples were washed two times with PBS-T for 5 minutes, counterstained with 1 µg/ml DAPI (Thermo Fisher, 1:1000 dilution) for 10 minutes, washed two times with PBS-T, and mounted on glass slides with ProLong Gold antifade reagent (Molecular Probes) for imaging.

Mounted coverslips were imaged as 7–11 optical sections on a Leica STELLARIS 5 LIAchroic confocal laser-scanning microscope with a 63x 1.4 NA plan apochromat oil-immersion objective and the following acquisition parameters: 60.13 nm pixel size at 3x optical zoom; 1000 × 1000 pixels^2^ field of view; 475 ns pixel dwell time; 330 nm z step size; 1 Airy Unit pinhole for a 520 nm emission; line accuracy = 2; 405 nm laser power = 1.7%; 488 nm laser power = 10%; 561 nm laser power = 5%; 638 nm laser power = 0.1–1%; and all detectors in photon-counting mode. Image stacks were deconvolved with Leica LIGHTNING deconvolution software using default parameters and Prolong Gold as the immersion medium.

### CPC droplet simulations

CPC simulations were initialized with a +1-state vertical stripe of width *W* at the midpoint overlaid with a +1-state circle with radius R_IC_ in the center of the domain (*L*_*x*_ = *L*_*y*_ = 2 = 6.4µm, GridSize = 512). The rest of the domain was filled with the –1 state. Reasonable values of *R*_*IC*_ were selected from HeLa measurements of the inner centromere (10 values between 50–350nm), and *W* values ranging from 50–140 nm. Irregular width simulations were initialized with the same values for *R*_*IC*_; *W* was selected from a normal distribution (*µ* = 90 nm, *σ* = 10 nm) at each vertical position and initialized for five simulations with the random seed set to the simulation number (50 total). Visualizations in Figures 5, 6A, and 6B show the central 3.2 µm of the domain—lateral mesh points outside this range factor into the calculations but do not change from their initial conditions. All simulations were run with *dt* = 1.53e-6 for 0.04 or 0.4 *t*_*char*_ (~7 or 70 minutes), which heuristically gave stable and accurate simulations for the resolution *dx* = 1/512 as determined by comparisons between SAV and NMG. For NMusing numpy.random.choice G, tolerance = 1e-5 and solver_iter = 1e4.

### Phos-Tag immunoblotting

Cells were synchronized with thymidine then RO-3306 and released for 10 minutes (HeLa) or 70 minutes (MCF10A-5E) as described above. Cells were counted using an automated cell counter (Invitrogen), lysed with Laemmli sample buffer (62.5 mM Tris-HCl (pH 6.8), 2% SDS, 10% glycerol, 100 mM dithiothreitol, and 0.01% bromophenol blue), and sheared with a 25-gauge needle (BD PrecisionGlide). Phos-tag immunoblotting of 50,000 cells per lane was performed on polyacrylamide gels of 6% (INCENP), 8% (AURKB and CDCA8), or 15% (BIRC5) containing 10 mM Phos-tag acrylamide (Nard Institute) and 0.1 mM MnO_4_•4H_2_O. Gels were run with WIDE-VIEW prestained protein marker (FUJIFILM Wako) under 40 mA constant current for 70–90 minutes. Before electrophoretic transfer, the gels were incubated in transfer buffer (25 mM Tris, 192 mM glycine, 0.0375% SDS) plus 1 mM EDTA for 10 minutes, followed by an additional 10-minute incubation in transfer buffer. Proteins were tank transferred to a PVDF membrane (Millipore) in transfer buffer plus 10% methanol (INCENP, AURKB, and CDCA8) or 20% methanol (BIRC5) at 100 V for 60 minutes on ice. Membrane blocking and antibody dilution were performed with 5% low-fat milk powder in Tris-buffered saline (TBS) containing 0.1% Tween 20 solution (TBS-T). Immunoblotting and secondary fluorescence or chemiluminescence detection was completed as described^68^ with primary antibodies for AURKB (BD Biosciences, 1:1000 dilution), BIRC5 (Cell Signaling, 1:1000 dilution), CDCA8 (P.T.S. custom rabbit polyclonal,^69^ 1:2000 dilution), GAPDH (Ambion, 1:10,000 dilution), INCENP (Cell Signaling, 1:1000 dilution), and vinculin (Millipore, 1:10,000 dilution).

## QUANTIFICATION AND STATISTICAL ANALYSIS

### Defining *R*_*i*_ and *R*_*eq*_

For critical droplet simulations, *R*(*t*) was defined at every 10 time steps with contourc.m and levels = 0. *R*_*i*_ was interpolated as the *R*_*0*_ halfway between the largest R_0_ that dissipated and the smallest *R*_*0*_ that persisted for a given *ϵ*_*m*_. *R*_*eq*_ was gleaned from the inflection point of *R*(*t*) for droplets that still dissipated at a given *ϵ*_*m*_. We smoothed numerical fluctuations in *R*(*t*) by fitting a fifth-order polynomial to the transition zone of interest and calculating zeros of the second derivative analytically. R_eq_ was regressed against *ϵ*_*m*_ with scipy.optimize.curve_fit using linear, hyperbolic, logarithmic, and hyperbolic-to-linear relationships. Alternative models were compared to the hyperbolic-to-linear fit by Bayes information criterion weights.^70^

### Quantification of CPC immunoreactive foci

Optical sections that transected spread chromosomes were maximum-intensity projected, and AURKB foci between sister chromatids were selected with the magic wand tool in ImageJ (tolerance = 40 [HeLa] and 25 with 8-connected mode [MCF10A-5E]). Segmented areas were converted to radii of equivalent circles for downstream analysis. Line scans were performed with the freehand line tool in ImageJ (width = 5 pixels), and peaks in the exported traces were identified with the Python function

‘scipy.signal.find_peaks’ using intensity values rescaled by max intensity with prominence = 15 (rescaled by max intensity), min_width = 100 nm and max_width = 800 nm.100 bootstraps of 50% subsampled images (*N =* 10 for Hela and *N* = 38 for MCF10A-5E) were generated using numpy.random.choice (Python) without replacement. Half of the subsampled images were used to constrain *ϵ*_*m*_ from *R*_*eq*_ (Figures 4D and S6B), and the remaining half were used for line scan analysis (Figures 6E, 6H, and S6D).

### Drop-to-drop separation calculations from simulations

For each irregular-width simulation, we identified the center of droplets by calculating the mean location of each contour with contourc.m (level = 0) and tracked each droplet throughout the simulation. Time points within each simulation were randomly sampled (without replacement) so that the number of timepoints x the number of simulations approximated the number of chromosomes for each cell line.

Each time point was then randomly paired with a chromosome arm length measured in the cell line of interest (Figure S4A), and the distances between droplet centers were calculated for any droplets vertically within the paired arm-length distance from the inner centromere.

### Comparison of drop-to-drop separation between simulations and experiments

Using the bootstrapped distributions of drop-to-drop distances from experiments, we binned the mean drop-to-drop distance as a histogram and estimated a 95% confidence interval. The optimal bin width was determined by comparing the 95% intervals to the average Gaussian kernel density estimate (using scipy.gaussian_kde) from these experimental bootstraps, which gave ~55 bins over the range 0– 3.5 µm.

### Phos-Tag densitometry

Immunoblots were quantified in FIJI as previously described.^68^ The intensity minimum between peaks was used to separate the upper and lower forms of CDCA8 and INCENP.

## REFERENCES

1. Laghmach, R., and Potoyan, D.A. (2021). Liquid-liquid phase separation driven compartmentalization of reactive nucleoplasm. Phys. Biol. 18, 015001. 10.1088/1478-3975/abc5ad.

2. Ibata, N., and Terentjev, E.M. (2022). Nucleation of cadherin clusters on cell-cell interfaces. Sci. Rep. 12, 18485. 10.1038/s41598-022-23220-x.

3. Yan, H., Konstorum, A., and Lowengrub, J.S. (2018). Three-Dimensional Spatiotemporal Modeling of Colon Cancer Organoids Reveals that Multimodal Control of Stem Cell Self-Renewal is a Critical Determinant of Size and Shape in Early Stages of Tumor Growth. Bull. Math. Biol. 80, 1404–1433. 10.1007/s11538-017-0294-1.

4. Wirtz, D., Du, W., Zhu, J., Wu, Y., Kiemen, A., Wan, Z., Hanna, E., and Sun, S. (2023). Mechano-induced homotypic patterned domain formation by monocytes. Res Sq. 10.21203/rs.3.rs-3372987/v1.

5. Yin, S., and Mahadevan, L. (2023). Contractility-Induced Phase Separation in Active Solids. Phys. Rev. Lett. 131, 148401. 10.1103/PhysRevLett.131.148401.

6. Huycke, T.R., Hakkinen, T.J., Miyazaki, H., Srivastava, V., Barruet, E., McGinnis, C.S., Kalantari, A., Cornwall-Scoones, J., Vaka, D., Zhu, Q., et al. (2024). Patterning and folding of intestinal villi by active mesenchymal dewetting. Cell 187, 3072–3089 e3020. 10.1016/j.cell.2024.04.039.

7. Chen, Y., and Ferrell, J.E., Jr. (2021). C. elegans colony formation as a condensation phenomenon. Nat Commun 12, 4947. 10.1038/s41467-021-25244-9.

8. Wang, Y., Li, S., Mokbel, M., May, A.I., Liang, Z., Zeng, Y., Wang, W., Zhang, H., Yu, F., Sporbeck, K., et al. (2024). Biomolecular condensates mediate bending and scission of endosome membranes. Nature 634, 1204–1210. 10.1038/s41586-024-07990-0.

9. Liu, Q.X., Doelman, A., Rottschafer, V., de Jager, M., Herman, P.M., Rietkerk, M., and van de Koppel, J. (2013). Phase separation explains a new class of self-organized spatial patterns in ecological systems. Proc. Natl. Acad. Sci. U. S. A. 110, 11905–11910. 10.1073/pnas.1222339110.

10. Sabari, B.R., Dall’Agnese, A., and Young, R.A. (2020). Biomolecular Condensates in the Nucleus. Trends Biochem. Sci. 45, 961–977. 10.1016/j.tibs.2020.06.007.

11. Lipinski, W.P., Visser, B.S., Robu, I., Fakhree, M.A.A., Lindhoud, S., Claessens, M., and Spruijt, E. (2022). Biomolecular condensates can both accelerate and suppress aggregation of alpha-synuclein. Sci Adv 8, eabq6495. 10.1126/sciadv.abq6495.

12. Deviri, D., and Safran, S.A. (2021). Physical theory of biological noise buffering by multicomponent phase separation. Proc. Natl. Acad. Sci. U. S. A. 118. 10.1073/pnas.2100099118.

13. Weirich, K.L., Banerjee, S., Dasbiswas, K., Witten, T.A., Vaikuntanathan, S., and Gardel, M.L. (2017). Liquid behavior of cross-linked actin bundles. Proc. Natl. Acad. Sci. U. S. A. 114, 2131–2136. 10.1073/pnas.1616133114.

14. Setru, S.U., Gouveia, B., Alfaro-Aco, R., Shaevitz, J.W., Stone, H.A., and Petry, S. (2021). A hydrodynamic instability drives protein droplet formation on microtubules to nucleate branches. Nat Phys 17, 493–498. 10.1038/s41567-020-01141-8.

15. Heltberg, M.S., Lucchetti, A., Hsieh, F.S., Minh Nguyen, D.P., Chen, S.H., and Jensen, M.H. (2022). Enhanced DNA repair through droplet formation and p53 oscillations. Cell 185, 4394–4408 e4310. 10.1016/j.cell.2022.10.004.

16. Anderson, D.M., McFadden, G.B., and Wheeler, A.A. (1998). Diffuse-interface methods in fluid mechanics. Annual Review of Fluid Mechanics 30, 139–165. DOI 10.1146/annurev.fluid.30.1.139.

17. Cahn, J.W., and Hilliard, J.E. (1958). Free Energy of a Nonuniform System .1. Interfacial Free Energy. J. Chem. Phys. 28, 258–267. Doi 10.1063/1.1744102.

18. Lee, C.Y., Jeong, D., Yang, J.X., and Kim, J. (2020). Nonlinear Multigrid Implementation for the Two-Dimensional Cahn-Hilliard Equation. Mathematics-Basel 8. ARTN 97 10.3390/math8010097.

19. Soares, E.d.A. Jr. A.G.B., and Tavares, F.W. (2023). Exponential Integrators for Phase-Field Equations using Pseudo-spectral Methods: A Python Implementation. arXiv, 2305.08998. https://arxiv.org/abs/2305.08998.

20. Langtangen, H.P., and Logg, A. (2016). Solving PDEs in Python : The FEniCS Tutorial I. Simula SpringerBriefs on Computing 3. 1st ed. Springer International Publishing : Imprint: Springer,.

21. Gouveia, B., Kim, Y., Shaevitz, J.W., Petry, S., Stone, H.A., and Brangwynne, C.P. (2022). Capillary forces generated by biomolecular condensates. Nature 609, 255–264. 10.1038/s41586-022-05138-6.

22. Brangwynne, C.P., Tompa, P., and Pappu, R.V. (2015). Polymer physics of intracellular phase transitions. Nature Physics 11, 899–904. 10.1038/Nphys3532.

23. Cain, J.Y., Yu, J.S., and Bagheri, N. (2023). The in silico lab: Improving academic code using lessons from biology. Cell Syst 14, 1–6. 10.1016/j.cels.2022.11.006.

24. Trivedi, P., Palomba, F., Niedzialkowska, E., Digman, M.A., Gratton, E., and Stukenberg, P.T. (2019). The inner centromere is a biomolecular condensate scaffolded by the chromosomal passenger complex. Nat. Cell Biol. 21, 1127–1137. 10.1038/s41556-019-0376-4.

25. Hedtfeld, M., Dammers, A., Koerner, C., and Musacchio, A. (2024). A validation strategy to assess the role of phase separation as a determinant of macromolecular localization. Mol. Cell 84, 1783–1801 e1787. 10.1016/j.molcel.2024.03.022.

26. Novickcohen, A., and Segel, L.A. (1984). Nonlinear Aspects of the Cahn-Hilliard Equation. Physica D 10, 277–298. Doi 10.1016/0167-2789(84)90180-5.

27. Choi, J.W., Lee, H.G., Jeong, D., and Kim, J. (2009). An unconditionally gradient stable numerical method for solving the Allen-Cahn equation. Physica A 388, 1791–1803. 10.1016/j.physa.2009.01.026.

28. Oliveira, G.M., Oravecz, A., Kobi, D., Maroquenne, M., Bystricky, K., Sexton, T., and Molina, N. (2021). Precise measurements of chromatin diffusion dynamics by modeling using Gaussian processes. Nat Commun 12, 6184. 10.1038/s41467-021-26466-7.

29. Matsuda, T., Miyawaki, A., and Nagai, T. (2008). Direct measurement of protein dynamics inside cells using a rationally designed photoconvertible protein. Nat. Methods 5, 339–345. 10.1038/nmeth.1193.

30. Trottenberg, U., Oosterlee, C.W., and Schüller, A. (2001). Multigrid (Academic Press).

31. Kim, J., Kang, K.K., and Lowengrub, J. (2004). Conservative multigrid methods for Cahn-Hilliard fluids. J Comput Phys 193, 511–543. 10.1016/j.jcp.2003.07.035.

32. Shen, J., Xu, J., and Yang, J. (2018). The scalar auxiliary variable (SAV) approach for gradient flows. J Comput Phys 353, 407–416. 10.1016/j.jcp.2017.10.021.

33. Frigo, M., and Johnson, S.G. (2005). The Design and Implementation of FFTW3. Proceedings of the IEEE 93, 216–231. 10.1109/JPROC.2004.840301.

34. Jiang, M.S., Zhang, Z.Y., and Zhao, J. (2022). Improving the accuracy and consistency of the scalar auxiliary variable (SAV) method with relaxation. J Comput Phys 456. ARTN 110954 10.1016/j.jcp.2022.110954.

35. Smith, A.E., Slepchenko, B.M., Schaff, J.C., Loew, L.M., and Macara, I.G. (2002). Systems analysis of Ran transport. Science 295, 488–491.

36. Wang, L., Paudel, B.B., McKnight, R.A., and Janes, K.A. (2023). Nucleocytoplasmic transport of active HER2 causes fractional escape from the DCIS-like state. Nat Commun 14, 2110. 10.1038/s41467-023-37914-x.

37. Albeck, J.G., Burke, J.M., Spencer, S.L., Lauffenburger, D.A., and Sorger, P.K. (2008). Modeling a snap-action, variable-delay switch controlling extrinsic cell death. PLoS Biol. 6, 2831–2852. 07-PLBI-RA-3412 [pii] 10.1371/journal.pbio.0060299.

38. Schoeberl, B., Eichler-Jonsson, C., Gilles, E.D., and Muller, G. (2002). Computational modeling of the dynamics of the MAP kinase cascade activated by surface and internalized EGF receptors. Nat. Biotechnol. 20, 370–375.

39. Shin, Y., and Brangwynne, C.P. (2017). Liquid phase condensation in cell physiology and disease. Science 357. 10.1126/science.aaf4382.

40. Lee, D., Huh, J.Y., Jeong, D., Shin, J., Yun, A., and Kim, J. (2014). Physical, mathematical, and numerical derivations of the Cahn-Hilliard equation. Computational Materials Science 81, 216–225. 10.1016/j.commatsci.2013.08.027.

41. Harris, C.R., Millman, K.J., van der Walt, S.J., Gommers, R., Virtanen, P., Cournapeau, D., Wieser, E., Taylor, J., Berg, S., Smith, N.J., et al. (2020). Array programming with NumPy. Nature 585, 357–362. 10.1038/s41586-020-2649-2.

42. Roesch, E., Greener, J.G., MacLean, A.L., Nassar, H., Rackauckas, C., Holy, T.E., and Stumpf, M.P.H. (2023). Julia for biologists. Nat. Methods 20, 655–664. 10.1038/s41592-023-01832-z.

43. Lin, C.W., Nocka, L.M., Stinger, B.L., DeGrandchamp, J.B., Lew, L.J.N., Alvarez, S., Phan, H.T., Kondo, Y., Kuriyan, J., and Groves, J.T. (2022). A two-component protein condensate of the EGFR cytoplasmic tail and Grb2 regulates Ras activation by SOS at the membrane. Proc. Natl. Acad. Sci. U. S. A. 119, e2122531119. 10.1073/pnas.2122531119.

44. Yue, P., Zhou, C., and Feng, J.J. (2007). Spontaneous shrinkage of drops and mass conservation in phase-field simulations. J Comput Phys 223, 1–9. 10.1016/j.jcp.2006.11.020.

45. Sweatt, A.J., Griffiths, C.D., Groves, S.M., Paudel, B.B., Wang, L., Kashatus, D.F., and Janes, K.A. (2024). Proteome-wide copy-number estimation from transcriptomics. Mol. Syst. Biol. 20, 1230–1256. 10.1038/s44320-024-00064-3.

46. Trivedi, P., and Stukenberg, P.T. (2016). A Centromere-Signaling Network Underlies the Coordination among Mitotic Events. Trends Biochem. Sci. 41, 160–174. 10.1016/j.tibs.2015.11.002.

47. Abad, M.A., Ruppert, J.G., Buzuk, L., Wear, M., Zou, J., Webb, K.M., Kelly, D.A., Voigt, P., Rappsilber, J., Earnshaw, W.C., and Jeyaprakash, A.A. (2019). Borealin-nucleosome interaction secures chromosome association of the chromosomal passenger complex. J. Cell Biol. 218, 3912–3925. 10.1083/jcb.201905040.

48. Yamagishi, Y., Honda, T., Tanno, Y., and Watanabe, Y. (2010). Two histone marks establish the inner centromere and chromosome bi-orientation. Science 330, 239–243. 10.1126/science.1194498.

49. Liang, J., Niu, Z., Zhang, B., Yu, X., Zheng, Y., Wang, C., Ren, H., Wang, M., Ruan, B., Qin, H., et al. (2021). p53-dependent elimination of aneuploid mitotic offspring by entosis. Cell Death Differ 28, 799–813. 10.1038/s41418-020-00645-3.

50. Samejima, K., Platani, M., Wolny, M., Ogawa, H., Vargiu, G., Knight, P.J., Peckham, M., and Earnshaw, W.C. (2015). The Inner Centromere Protein (INCENP) Coil Is a Single alpha-Helix (SAH) Domain That Binds Directly to Microtubules and Is Important for Chromosome Passenger Complex (CPC) Localization and Function in Mitosis. J. Biol. Chem. 290, 21460–21472. 10.1074/jbc.M115.645317.

51. Banani, S.F., Lee, H.O., Hyman, A.A., and Rosen, M.K. (2017). Biomolecular condensates: organizers of cellular biochemistry. Nat. Rev. Mol. Cell Biol. 18, 285–298. 10.1038/nrm.2017.7.

52. Anderson, D.E., Losada, A., Erickson, H.P., and Hirano, T. (2002). Condensin and cohesin display different arm conformations with characteristic hinge angles. J. Cell Biol. 156, 419–424. 10.1083/jcb.200111002.

53. Zhou, L., Liang, C., Chen, Q., Zhang, Z., Zhang, B., Yan, H., Qi, F., Zhang, M., Yi, Q., Guan, Y., et al. (2017). The N-Terminal Non-Kinase-Domain-Mediated Binding of Haspin to Pds5B Protects Centromeric Cohesion in Mitosis. Curr. Biol. 27, 992–1004. 10.1016/j.cub.2017.02.019.

54. Villa, F., Capasso, P., Tortorici, M., Forneris, F., de Marco, A., Mattevi, A., and Musacchio, A. (2009). Crystal structure of the catalytic domain of Haspin, an atypical kinase implicated in chromatin organization. Proc. Natl. Acad. Sci. U. S. A. 106, 20204–20209. 10.1073/pnas.0908485106.

55. Kohn, J.E., Millett, I.S., Jacob, J., Zagrovic, B., Dillon, T.M., Cingel, N., Dothager, R.S., Seifert, S., Thiyagarajan, P., Sosnick, T.R., et al. (2004). Random-coil behavior and the dimensions of chemically unfolded proteins. Proc. Natl. Acad. Sci. U. S. A. 101, 12491–12496. 10.1073/pnas.0403643101.

56. Gascoigne, K.E., Takeuchi, K., Suzuki, A., Hori, T., Fukagawa, T., and Cheeseman, I.M. (2011). Induced ectopic kinetochore assembly bypasses the requirement for CENP-A nucleosomes. Cell 145, 410–422. 10.1016/j.cell.2011.03.031.

57. Broad, A.J., DeLuca, K.F., and DeLuca, J.G. (2020). Aurora B kinase is recruited to multiple discrete kinetochore and centromere regions in human cells. J. Cell Biol. 219. 10.1083/jcb.201905144.

58. Liu, Y., Mi, Y., Mueller, T., Kreibich, S., Williams, E.G., Van Drogen, A., Borel, C., Frank, M., Germain, P.L., Bludau, I., et al. (2019). Multi-omic measurements of heterogeneity in HeLa cells across laboratories. Nat. Biotechnol. 37, 314–322. 10.1038/s41587-019-0037-y.

59. Janes, K.A., Wang, C.C., Holmberg, K.J., Cabral, K., and Brugge, J.S. (2010). Identifying single-cell molecular programs by stochastic profiling. Nat. Methods 7, 311–317. 10.1038/nmeth.1442.

60. Soule, H.D., Maloney, T.M., Wolman, S.R., Peterson, W.D., Jr., Brenz, R., McGrath, C.M., Russo, J., Pauley, R.J., Jones, R.F., and Brooks, S.C. (1990). Isolation and characterization of a spontaneously immortalized human breast epithelial cell line, MCF-10. Cancer Res. 50, 6075–6086.

61. Martin, I.M., Aponte-Santamaria, C., Schmidt, L., Hedtfeld, M., Iusupov, A., Musacchio, A., and Grater, F. (2022). Phosphorylation tunes elongation propensity and cohesiveness of INCENP’s intrinsically disordered region. J. Mol. Biol. 434, 167387. 10.1016/j.jmb.2021.167387.

62. Rai, A.K., Chen, J.X., Selbach, M., and Pelkmans, L. (2018). Kinase-controlled phase transition of membraneless organelles in mitosis. Nature 559, 211–216. 10.1038/s41586-018-0279-8.

63. Riback, J.A., Eeftens, J.M., Lee, D.S.W., Quinodoz, S.A., Donlic, A., Orlovsky, N., Wiesner, L., Beckers, L., Becker, L.A., Strom, A.R., et al. (2023). Viscoelasticity and advective flow of RNA underlies nucleolar form and function. Mol. Cell 83, 3095–3107 e3099. 10.1016/j.molcel.2023.08.006.

64. Lee, A.A., Kim, N.H., Alvarez, S., Ren, H., DeGrandchamp, J.B., Lew, L.J.N., and Groves, J.T. (2024). Bimodality in Ras signaling originates from processivity of the Ras activator SOS without deterministic bistability. Sci Adv 10, eadi0707. 10.1126/sciadv.adi0707.

65. Soding, J., Zwicker, D., Sohrabi-Jahromi, S., Boehning, M., and Kirschbaum, J. (2020). Mechanisms for Active Regulation of Biomolecular Condensates. Trends Cell Biol. 30, 4–14. 10.1016/j.tcb.2019.10.006.

66. Bretin, E., Denis, R., Masnou, S., Sengers, A., and Terii, G. (2023). A multiphase Cahn-Hilliard system with mobilities and the numerical simulation of dewetting. Esaim-Mathematical Modelling and Numerical Analysis 57, 1473–1509. 10.1051/m2an/2023023.

67. Chen, L., Zhao, J., and Yang, X. (2018). Regularized linear schemes for the molecular beam epitaxy model with slope selection. Applied Numerical Mathematics 128, 139–156. 10.1016/j.apnum.2018.02.004.

68. Janes, K.A. (2015). An analysis of critical factors for quantitative immunoblotting. Sci Signal 8, rs2. 10.1126/scisignal.2005966.

69. Trivedi, P., Zaytsev, A.V., Godzi, M., Ataullakhanov, F.I., Grishchuk, E.L., and Stukenberg, P.T. (2019). The binding of Borealin to microtubules underlies a tension independent kinetochore-microtubule error correction pathway. Nat Commun 10, 682. 10.1038/s41467-019-08418-4.

70. Wagenmakers, E.J., and Farrell, S. (2004). AIC model selection using Akaike weights. Psychon Bull Rev 11, 192–196. 10.3758/bf03206482.

