## Supplemental Data 1 for "Cahn-Hilliard dynamical models for condensed biomolecular systems"

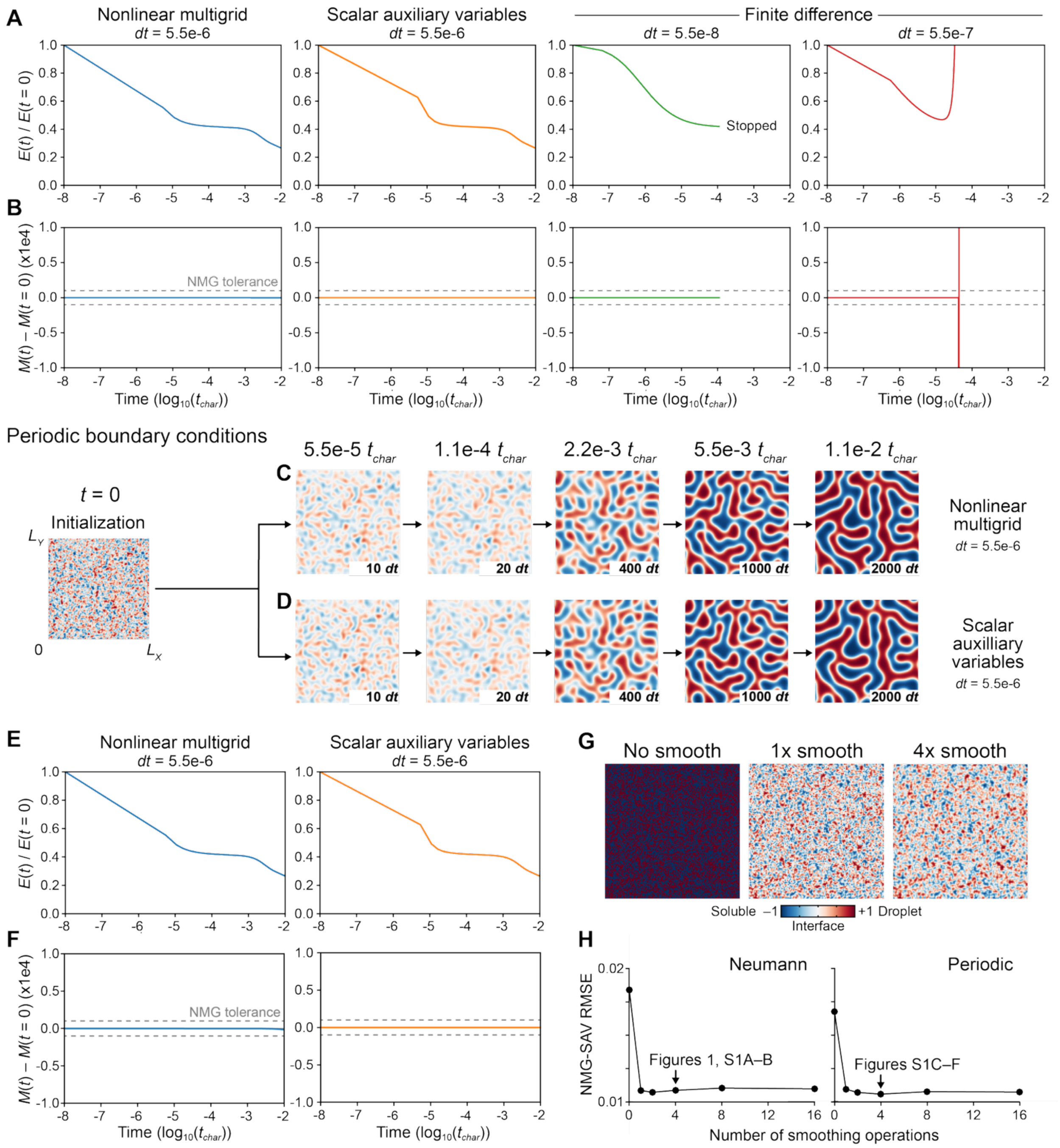


**Figure S1. Extended spinodal decompositions comparing different solvers. Related to Figure 1.**
(A and B) Time evolution for the relative energy [$E(t)$ / $E(t = 0)$] (A) and mass error [$M(t) – M(t = 0)$] (B) of the spinodal decompositions in Figure 1.
(C and D) Time evolution for the same spinodal decomposition as in Figure 1 but with periodic boundary conditions.
(E and F) Time evolution for the relative energy (E) and mass error (F) of the spinodal decompositions in (C) and (D).
(G) A 50:50 initialization of ±1 chemical states with zero (left), two (center), or four (right) rounds of smoothing.
(H) Energy minimizations of spinodal decomposition are consistent and stable after at-least one round of smoothing. Root mean squared error (RMSE) is shown as a function of the number of smoothing operations. The level of smoothing used in the indicated figures is highlighted.

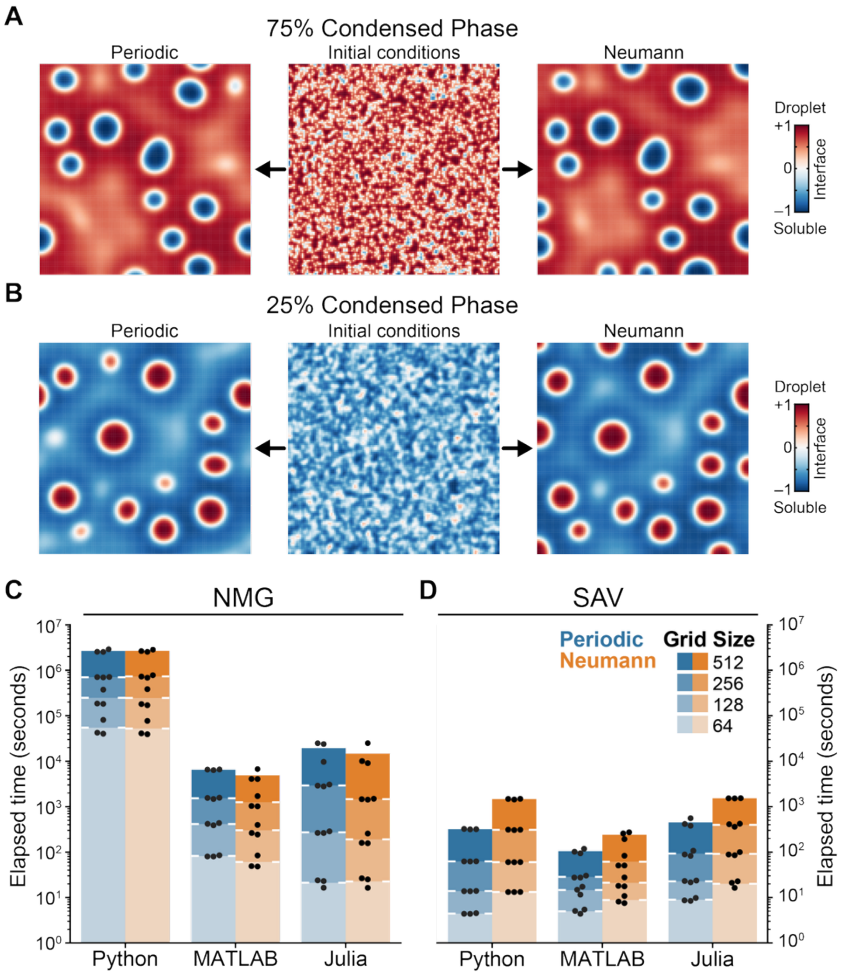


**Figure S2. Alternative initial conditions for spinodal decomposition, NMG–SAV error, and performance for different mesh sizes. Related to Figure 2.**

(A and B) Snapshots of spinodal decomposition for initial conditions with 75% +1 state (A) or 25% +1 state (B) and periodic (left) or Neumann (right) boundary conditions.

(C and D) Runtime performance across mesh sizes ($2^5$–$2^8$) for NMG (C) and SAV (D) solvers in Python, MATLAB, and Julia for spinodal decompositions initialized with $N$ = 3 random mixtures of ±1 chemical states (25:75, 50:50, and 75:25) and either periodic or Neumann boundary conditions. All simulations were performed on a Linux x86_64 processor with 100 GB RAM on a $2^7$ x $2^7$ mesh ($L_X = L_Y = 1$) for 2000 time steps ($dt$ = 5.5e-6) and $\epsilon_{m=8}$.

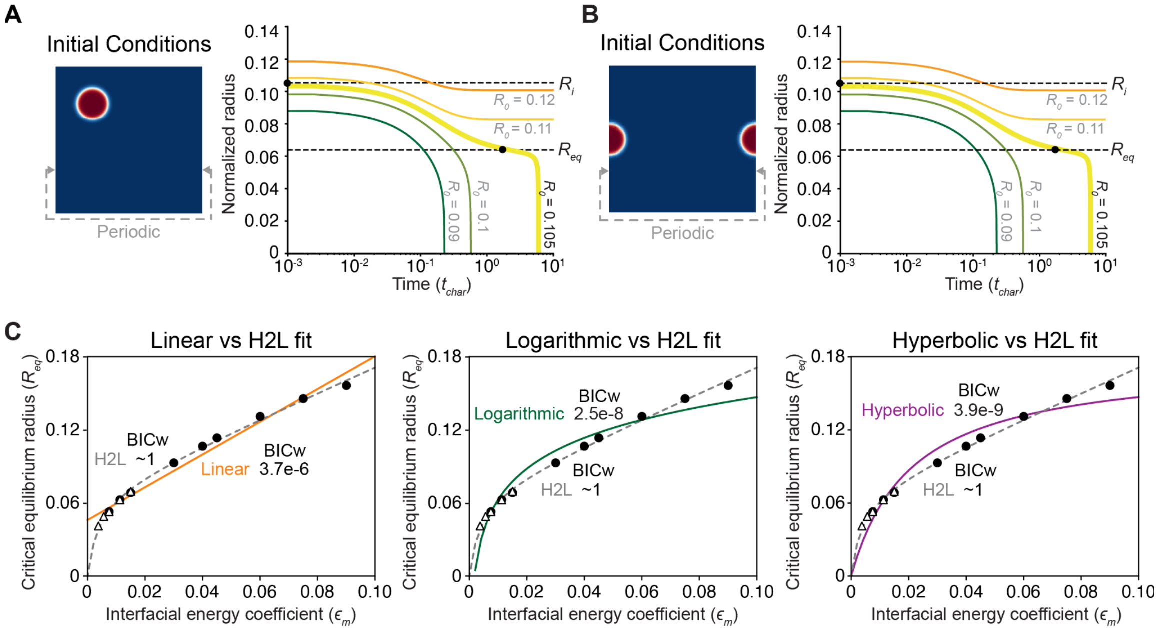


**Figure S3. Robustness of critical radii and fit of $R_{eq}$ compared to alternatives. Related to Figure 3.**
(A and B) Identical simulations as in Figure 3B but with an off-center droplet (A) or a droplet split across a periodic boundary (B). $R_0$ = 0.105 is shown as an example.
(C) The hyperbolic-to-linear (H2L) fit of the data in Figure 3E versus a linear (left), logarithmic (middle), or hyperbolic (right) model. Fits were compared by Bayes information criterion (BIC), and BIC weights[1] (BICw) are shown indicating the relative likelihood of each alternative compared to H2L.

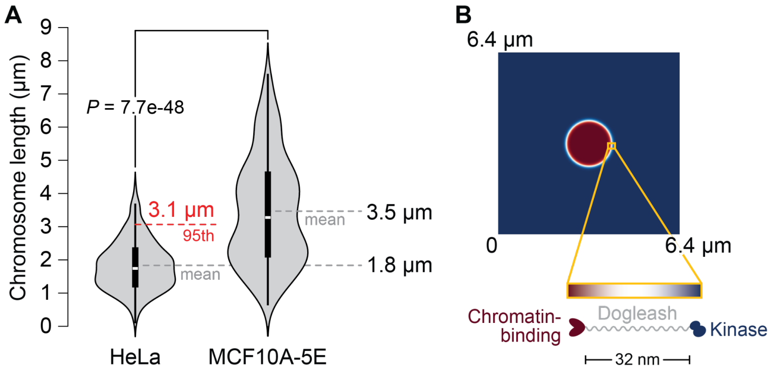


**Figure S4. Mapping the Cahn-Hilliard equation to physical dimensions of (pro)metaphase chromosomes and the CPC. Related to Figure 4.**

(A) Size distribution of spread chromosomes prepared from HeLa cells and MCF10A-5E cells. Data are from *N* = 231 chromosomes from 10 images for HeLa and *N* = 218 chromosomes from 38 images for MCF10A-5E. Distributions were compared by KS test.

(B) Plausibility of the alpha-helical “dogleash” subunit of the CPC spanning condensed and soluble phases. Given a 3.2 µm arm length (A) x 2 arms = 6.4 µm domain, $\epsilon_m$ = 21.6 ± 0.6 nm for HeLa and $\epsilon_m$ = 28.5 nm ± 1.1 nm for MCF10A-5E (Equation 8).

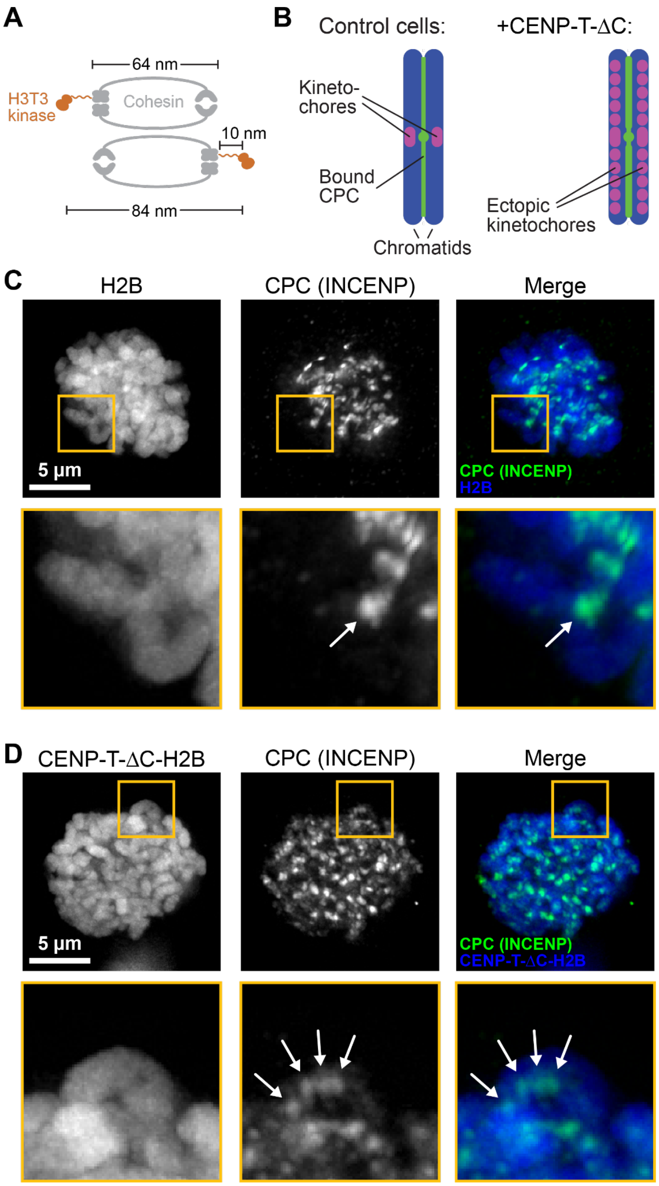


**Figure S5. Estimation and perturbation of the pH3T3 width. Related to Figure 5.**
(A) Putative configuration of the cohesin complex with H3T3 kinase. The cohesin ring holds together sister chromatids at different orientations, and one of its regulatory subunits binds the unstructured N terminus of H3T3 kinase.[2,3] Length estimates are from Ref. [4,5].
(B) Kinetochore perturbation described by Gascoigne et al.[6] HeLa cells transiently coexpressing GFP-CENP-C-ΔC-H2B and mCherry-CENP-T-ΔC-H2B (not shown) were compared to control cells transiently coexpressing GFP-H2B and mCherry-H2B.
(C and D) Enlarged and reprocessed images from Figure S3C of Ref. [6]. Reprinted with permission from Cell Press.

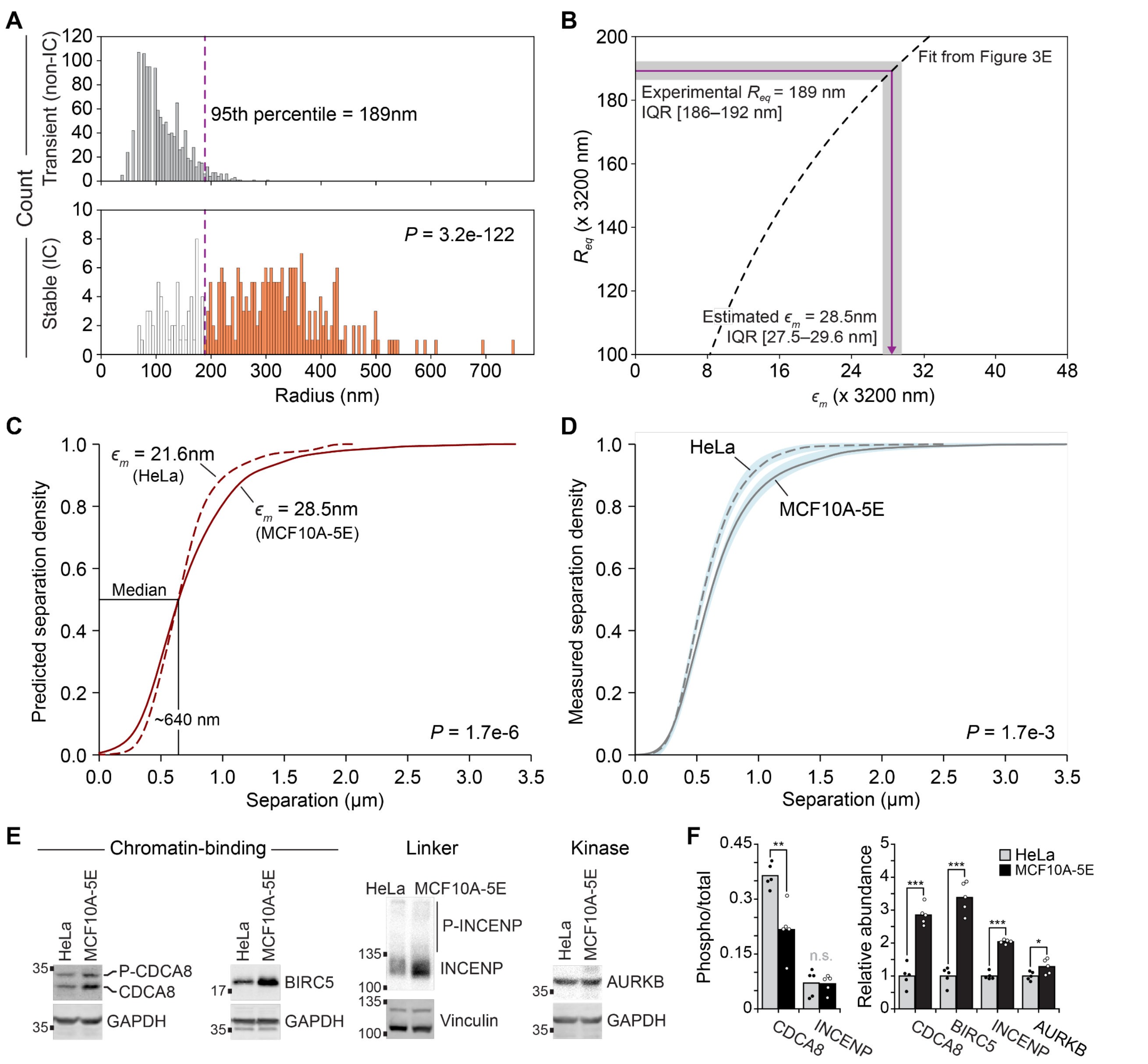


**Figure S6. HeLa- and MCF10A-5E-specific differences in CPC properties. Related to Figure 6.**
(A) Histogram of non-IC foci (gray) and IC foci (white and orange) quantified by idealized circular radius from $N = 218$ chromosomes in 50 MCF10A cells. The 95th percentile of the non-IC droplet size distribution defining $R_{eq}$ is shown. The distribution of non-IC foci ($N = 1223$) and IC foci ($N = 268$) were compared by KS test.
(B) Estimation of $\epsilon_m$ from measured $R_{eq}$ using the hyperbolic-to-linear regression (Figure 3E) scaled for a 3200-nm physical spatial domain. The interquartile range (IQR) was calculated by 50% subsampling of $N = 38$ MCF10A-5E cells for 100 iterations without replacement and propagating to the $\epsilon_m$ estimate.
(C) Cumulative density function (CDF) plot of drop-to-drop distances from Cahn-Hilliard simulations with $\epsilon_m = 21.6$ nm (dotted line, $N = 1221$ distances) or 28.5 nm (solid line, $N = 839$ distances). The distributions were compared by KS test.
(D) Median CDF plot ± 95% bootstrapped confidence interval (blue) of peak-to-peak separation for CPC foci in HeLa cells (dotted line, $N = 583$ peaks) or MCF10A cells (solid line, $N = 831$ peaks). The distributions were compared by KS test.
(E) Representative Phos-Tag immunoblots of whole-cell extracts (50,000 cells) from HeLa and MCF10A-5E cells. Lysates were immunoblotted for the indicated targets with GAPDH or vinculin used as loading controls.

(F) Quantitative densitometry of (E) for $N = 5$ biological replicates of each cell line. *$P < 0.05$, **$P < 0.01$, ***$P < 0.0001$ by unpaired two-sample $t$ test.

### SUPPLEMENTARY REFERENCES


1. Wagenmakers, E.J., and Farrell, S. (2004). AIC model selection using Akaike weights. Psychon Bull Rev *11*, 192-196. 10.3758/bf03206482.
2. Zhou, L., Liang, C., Chen, Q., Zhang, Z., Zhang, B., Yan, H., Qi, F., Zhang, M., Yi, Q., Guan, Y., et al. (2017). The N-Terminal Non-Kinase-Domain-Mediated Binding of Haspin to Pds5B Protects Centromeric Cohesion in Mitosis. Curr. Biol. *27*, 992-1004. 10.1016/j.cub.2017.02.019.
3. Villa, F., Capasso, P., Tortorici, M., Forneris, F., de Marco, A., Mattevi, A., and Musacchio, A. (2009). Crystal structure of the catalytic domain of Haspin, an atypical kinase implicated in chromatin organization. Proc. Natl. Acad. Sci. U. S. A. *106*, 20204-20209. 10.1073/pnas.0908485106.
4. Anderson, D.E., Losada, A., Erickson, H.P., and Hirano, T. (2002). Condensin and cohesin display different arm conformations with characteristic hinge angles. J. Cell Biol. *156*, 419-424. 10.1083/jcb.200111002.
5. Kohn, J.E., Millett, I.S., Jacob, J., Zagrovic, B., Dillon, T.M., Cingel, N., Dothager, R.S., Seifert, S., Thiyagarajan, P., Sosnick, T.R., et al. (2004). Random-coil behavior and the dimensions of chemically unfolded proteins. Proc. Natl. Acad. Sci. U. S. A. *101*, 12491-12496. 10.1073/pnas.0403643101.
6. Gascoigne, K.E., Takeuchi, K., Suzuki, A., Hori, T., Fukagawa, T., and Cheeseman, I.M. (2011). Induced ectopic kinetochore assembly bypasses the requirement for CENP-A nucleosomes. Cell *145*, 410-422. 10.1016/j.cell.2011.03.031.
